# SNV-FEAST: microbial source tracking with single nucleotide variants

**DOI:** 10.1101/2022.05.28.493810

**Authors:** Leah Briscoe, Eran Halperin, Nandita R. Garud

## Abstract

Elucidating the sources of a microbiome can provide insight into the ecological dynamics responsible for the formation of these communities. “Source tracking” approaches to date leverage species abundance information, however, single nucleotide variants (SNVs) may be more informative because of their high specificity to certain sources. To overcome the computational burden of utilizing all SNVs for a given sample, we introduce a novel method to identify signature SNVs for source tracking. We show that signature SNVs used as input into a previously designed source tracking algorithm, FEAST, can more accurately estimate contributions than species and provide novel insights, demonstrated in three case studies.

## BACKGROUND

Understanding the sources that could contribute to the formation of a given microbiome is of great interest in elucidating the ecological processes that give rise to these complex communities and the impact of these communities on human and environmental health. For example, a hospital environment may introduce antibiotic resistance genes to an infant gut microbiome, and local selective pressures may result in vastly different microbial compositions in different parts of the ocean. Approaches for determining the proportion of a microbiome of interest (the “sink”) that is attributed to different microbiomes (the “sources”) is known as “source tracking” (Knights et al., 2011; Shenhav et al., 2019). Source tracking is useful for forensics, categorization of samples, detecting contamination, and tracing transmissions between different hosts or environments. While source tracking was developed as a way to quantitatively characterize a sample based on a set of samples with known origin, in most studies, the true source of samples may never be collected. In these cases, source tracking approaches are useful in identifying similarities between microbiome samples even if they cannot be used to definitively identify the true source of origin.

Current approaches for source tracking include the Bayesian approach, SourceTracker (Knights et al., 2011) and more recently the expectation-maximization approach, FEAST (Shenhav et al., 2019). These source tracking methods use species abundance profiles of the sample of interest (the sink) and of potential sources and compute percentages of sinks that are attributable to each potential source. However, species abundance profiles miss important subspecies single nucleotide variants (SNVs), which may provide higher resolution information than species about transmission patterns. For example, (Nayfach et al., 2016) found that the sharing of microbiome SNVs private to mothers and their infants decreases over the first year of the infant’s life while species sharing increases. This suggests that while the infant microbiome increasingly resembles the adult microbiome ecologically, sources other than the mother also colonize the infant. Thus, species-level resolution may obscure true sources of microbes while SNVs can reveal actual transmissions to the infant.

While tracking strain transmissions with SNVs has been highly successful in a number of studies (Asnicar et al., 2017a; Ferretti et al., 2018; Korpela et al., 2018; Li et al., 2016; Nayfach et al., 2016; Olm et al., 2021; Schmidt et al., 2019) current approaches to strain tracking are limited. These methods provide binary information by inferring whether or not a strain transmission has occurred per species but they do not shed light on the relative proportions of microbiomes that are similar. A specific example of this is inStrain (Olm et al., 2021) which computes a pairwise population-level average nucleotide identity (popANI) between two samples. If an infant harbors several strains derived from the mother at low frequency, these shared strains will have high popANI values, but they will represent a relatively small proportion of the infant’s microbiome. By contrast, source tracking allows us to simultaneously infer the putative proportions for multiple sources contributing to a given sink, integrated over all community members in the sink. As shown in **Figure 1**, one may be able to estimate that an infant microbiome is explained 25% by the mother, 10% by the dog, and 30% by unknown sources (Knights et al., 2011; Shenhav et al., 2019). In other words, source tracking with SNVs leverages not only the genetic variants within species, but also the relative abundances of the species that carry the SNVs.

**Figure 1:**
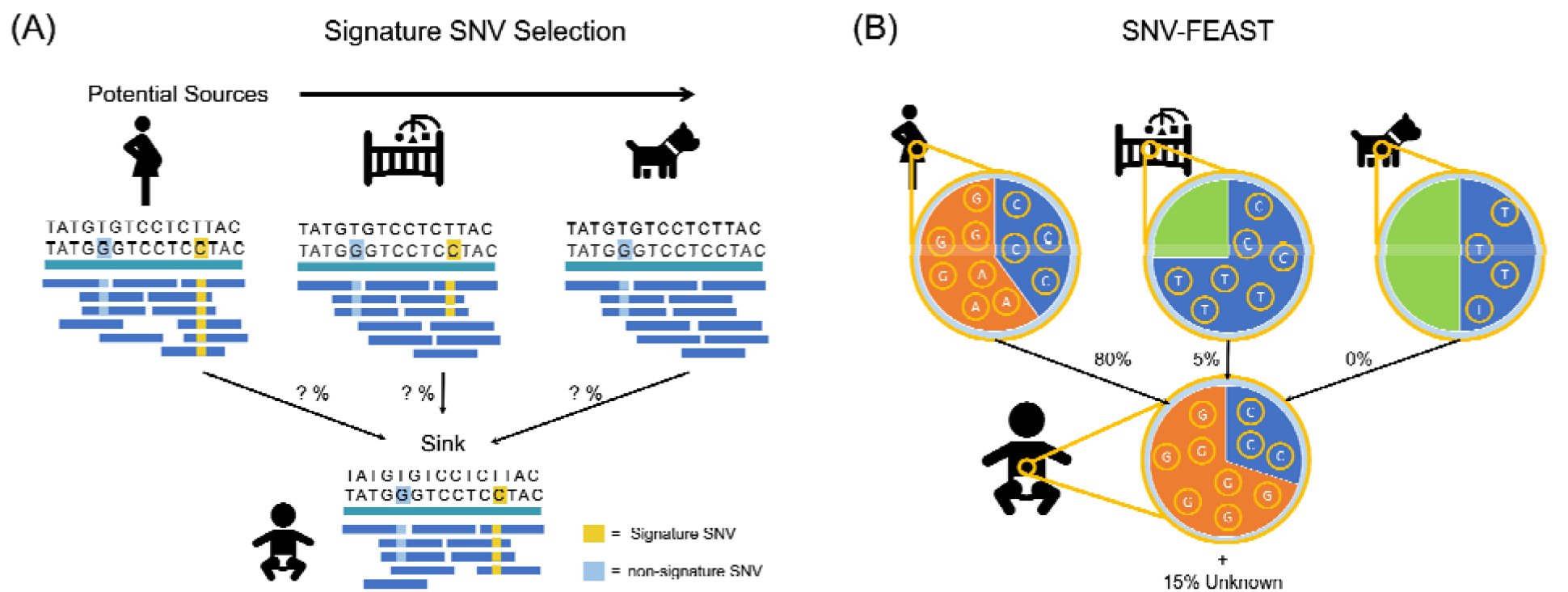
Signature SNV selection and SNV-FEAST. (A) A signature SNV is present in one or few but not all sources. By contrast, a non-signature SNV is generically present in multiple sources and thus provides little discriminating information. (B) SNV-FEAST estimates the proportion a given sink derived from various sources using the read counts for each allele in sinks and sources.

Here, we evaluate whether source contributions estimated with SNVs are more accurate than with only species when provided as inputs to FEAST (Shenhav et al., 2019) (hereafter referred to as SNV-FEAST and species-FEAST, respectively). FEAST (Shenhav et al., 2019) is faster and more accurate than previous source tracking tools (Knights et al., 2011), and therefore, is ideal for adaptation to SNV source tracking since it can accept larger numbers of features and input sources. Despite this improved computational efficiency, the potentially millions of single nucleotide variants across all microbiome species in a given host still can computationally overwhelm FEAST. To address this, we introduce a novel approach to determine signature SNVs that can be used as input to FEAST. This both reduces memory requirements and computation time in the FEAST estimation, allowing us to optimally estimate the source contribution of a sink. We find that SNV-FEAST and species-FEAST yield different outcomes when applied to simulated data, with SNV-FEAST frequently out-performing species-FEAST. We apply SNV-FEAST to three real-world case studies, including source tracking between infants and their mothers in the first year of life, between infants and the neonatal intensive care unit (NICU), and between oceans around the world. We confirm the ability of SNV-FEAST by recapitulating several previously published findings in our case studies, as well as discover new source tracking patterns across oceans. In sum, we show that SNVs can be used to estimate potential transmissions across hosts and across environments.

## RESULTS

### SNV-FEAST algorithm

Here we adapt FEAST to accept SNV abundance instead of species abundance as input. A computational challenge in using SNVs instead of species as input to FEAST is that SNVs contribute a significantly larger feature space. The number of different species comprising a microbiome can range from a few hundred to a few thousand, while the number of possible SNVs for a given species alone can be in the thousands (Schloissnig et al., 2013). This difference in number of input features can result in FEAST runtimes that last several hours instead of a few minutes and memory intensive storage of read counts at all sites of variation.

We devised a likelihood-based approach for selecting a set of informative or “signature” SNVs for a given source tracking analysis, allowing us to overcome the time and memory intensive challenges of utilizing SNV-level data. We identify these informative SNVs by computing a signature score (**Figure 1A**) (see **Methods)** that quantifies the extent to which SNVs in the sink are most likely derived from one of the potential sources. This is analogous to identifying SNVs private to sources and their sinks, but more generalized to include SNVs that may be found in multiple sources, albeit at higher frequency in one of the potential sources (see **Methods**).

To compute a signature score for a given SNV, two hypotheses are compared for each potential source: (1) that one source solely explains the observed allele counts in the sink and (2) all sources except that one source collectively explain the observed allele counts in the sink. For each hypothesis, we calculate the binomial log-likelihood for the estimate of the allele frequency in the sink, θ.

**Hypothesis 1:** Source *i* with allele frequency *p_i_* explains the allele counts in the sink.

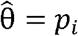

**Hypothesis 2:** A combination of all other sources except *i* (sources *j* ≠ *i*) explain the observed allele count distribution in the sink. The estimate of the sink allele frequency is computed using a mixture of the allele frequencies *p_j_* from those sources. The mixing parameter *α_j_* is learned using Sequential Least Squares Programming with the constraint that ∑_*j*≠*i*_ *α_j_* = 1.

The binomial log-likelihood is calculated as follows, where there are *n* reads with the reference allele and *m* reads with the alternative allele in the sink.

A log likelihood ratio representing the support for hypothesis 1 relative to hypothesis 2 is calculated per site per potential source. The maximum log likelihood ratio per site is the signature score for that SNV, representing how favorably one of the sources explains the sink over all other sources. Signature SNVs are those with scores greater than two standard deviations over the mean signature score computed for all SNVs (**Methods**).

### Evaluation of SNV-FEAST in simulations

To compare the accuracy of species-FEAST and SNV-FEAST, we performed simulations mimicking mother-infant transmissions with the goal of estimating contributions of different sources to an infant sink. Our simulations tested the ability of SNVs and species to recapitulate the true source composition in synthetic samples comprised of a mixture of reads drawn from multiple real fecal adult samples. To construct these synthetic infant microbiomes, we mixed metagenomic data from mothers sampled in a mother-infant dataset (Bäckhed et al., 2015) at various proportions as described below (**Methods**).

The difficulty of source tracking increases with the number of contributing sources (Shenhav et al., 2019). Thus, we simulate infants that have a small (<=5) versus large (6 - 10) number of contributing sources (**Supplementary Table 1**), including an unknown source (e.g. a randomly selected unrelated mother). Known source contributions to the simulated gut microbiome sample of the infant were varied between 1 and 90% while the unknown contribution varied between 10 and 90%. The unknown source was not presented to FEAST as a potential known source.

Additionally, not all species in a mother are transmitted to the infant (Asnicar et al., 2017b; Ferretti et al., 2018; Korpela et al., 2018; Sprockett et al., 2020; Yassour et al., 2018). Thus, in our simulations, species transmission rates were determined using a beta distribution, which is a natural model for values between (0,1) and often proposed for microbial abundance data (E. Z. Chen & Li, 2016; Martin et al., 2020; Sloan et al., 2006, 2007) (see **Methods**). We therefore consider four simulated scenarios: a combination of low versus high number of sources and low versus high transmission rates (see **Methods**).

**Figure 2** compares the performance of SNV-FEAST and species-FEAST in estimating the true contribution of sources. FEAST using SNVs has equal if not better performance than species in most scenarios, and performs especially well when transmission rates are low and unknown source proportions are high. SNVs have a lower root mean squared error (RMSE) compared to species in three of the four scenarios and higher Pearson correlation between true and estimated contributions in all four scenarios. The difference in these correlations for SNVs versus species is significant in all four cases when using a paired Wilcoxon signed rank test (high transmission: p-value = 0.00560, 0.00251 for small and large number of sources, low transmission: p-value = 0.00024, 0.002340 for small and large number of sources). These results suggest that SNVs may offer useful signatures of transmission.

**Figure 2:**
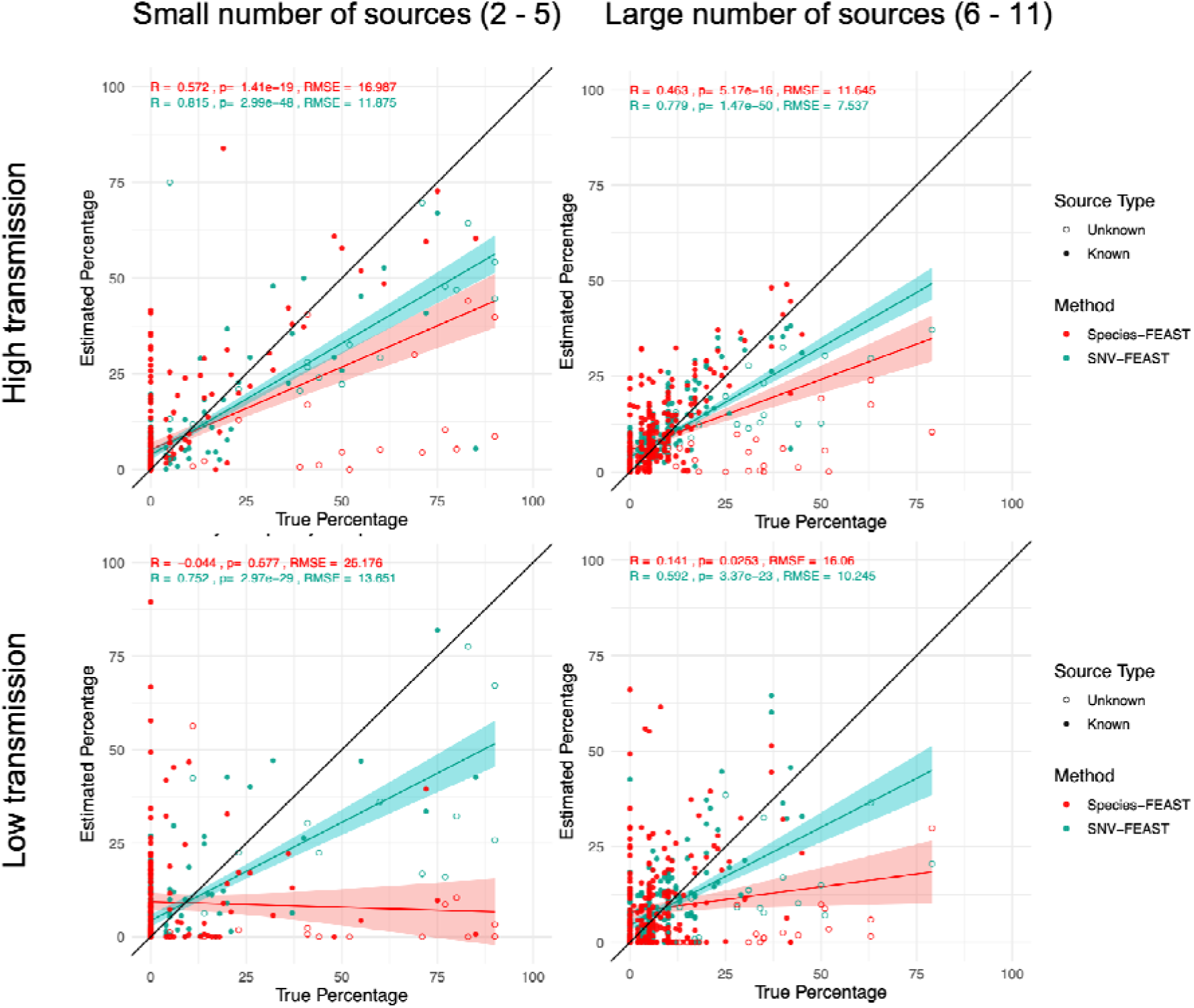
Ability of SNV and species-FEAST to recapitulate true contributions in simulations. Estimated known and unknown source proportions for infant microbiomes simulated with in silico mixtures of real maternal fecal microbiomes under different scenarios: either low number of contributing sources (<=5) or high number of sources (6-11), and high transmission rate of species or low transmission rate. Transmission rate is the probability of an infant being colonized by a given species, simulated using a beta distribution centered on the relative abundance of species in sources (**Methods**). 23 infants were simulated with five or fewer sources and 19 infants were simulated with a large number of sources (**Table S1**). The black line indicates the ground truth for proportions. For each simulated infant, there are 11 points plotted, whereby 10 correspond to known sources (some of which have zero contribution), and one corresponds to an unknown source which are indicated by a hollow circles in the plot.

To assess whether all species and all signatures SNVs in the sink are needed for accurate source tracking, we varied the proportion of species (from 10%, 50% or 100%) and SNVs (from 10%, 50% or 100%) included as inputs to the algorithm (**Figure S1**). We used Pearson correlation between the true and estimated proportions to represent accuracy of SNV-FEAST. When decreasing the percentage of SNVs used, there is no statistically significant change in the performance. However, when decreasing the percentage of species used, there are statistically significant decreases in the performance (**Figure S1**).

To illustrate the advantage of SNV-FEAST over traditional strain tracking approaches such as inStrain (Olm et al., 2021), we used the same synthetic communities produced in the above simulation for inStrain profiling between each infant and each of their potential contributing sources (**Figure S2**). InStrain computes a popANI score, which represents the average nucleotide identity between two different metagenomic samples for a given species. As per the inStrain paper, popANI values > 99.999% represent the same strain for that species being shared between samples (**Methods**). However, this approach provides a binarization as to whether or not a strain was transmitted, and does not account for the relative abundance of the strain in the sink. Thus, we computed the fraction of each infant’s species that have popANI ≥99.999%, with each potential source.

As expected, both SNV-FEAST and inStrain produce estimates of sharing that correlate positively with the ground truth mixture proportions of the contributing source samples in each infant (**Figure S2**). We found inStrain results yielded a 0.742 Pearson correlation (p < 1×10^-12^) with the true mixture proportions, whereas SNV-FEAST has a 0.866 Pearson correlation (p < 1×10^-12^) with the true proportions. The higher correlation values for SNV-FEAST likely reflect that relative abundances of strains and their genomic identities are simultaneously taken into account for source tracking, whereas inStrain only accounts for genomic identities. Finally, several of the shared species in the simulations had popANI values < 99.999%, reflecting the complex mixtures from multiple sources.

We next compared SNV-FEAST with the strain tracking procedure in Nayfach et al. 2016. Again, we used the same synthetic communities produced in the simulation to determine marker alleles as defined in Nayfach et al. 2016 (**Methods**). Here a marker allele is determined to be a SNV that is private to mother, infant, or the mother-infant dyad, and absent from the background population, which consisted of other samples in the dataset as well as samples from United States adults in the Human Microbiome Project (**Methods**). Species with ≥ 5% marker allele sharing between mother and infant were deemed to share a strain (**Methods**). We found a high correlation between the true mixture proportions (on x-axis) and the percentage of species with transmission events (y-axis) (Pearson correlation 0.915, p < 1 × 10^-16^). The higher correlation for the Nayfach et al. 2016 approach compared to the inStrain approach possibly reflects horizontal gene transfers between lineages residing in infants and mothers. By contrast, there was a lower correlation between the true mixture proportions (x-axis) and the sharing for all marker alleles across species present in the infant (y-axis) and (0.575 Pearson correlation, p < 1 × 10^-16^) (**Figure S3B**).

### Source tracking in infants over the first year of life

Having assessed the abilities of SNV-FEAST in synthetic data, we next estimated the contribution from the true mother over time to the true infant with SNV and species-FEAST in the Backhed et al. 2015 dataset. This dataset is composed of metagenomic samples from infants collected at four days, four months, and 12 months after birth, as well as their mothers at the time of delivery. Previous analyses on this data have shown that even while species similarity increases, infants and their mothers share fewer proportions of strains over time as revealed by sharing of SNVs private to mother-infant dyads (Nayfach et al., 2016). Thus, SNVs belonging to strains shared only by the infant and their mother may be more informative of the true source compared to species. Here we sought to test whether SNV and species-FEAST recapitulate these results (**Methods**).

In applying FEAST to the Backhed et al. 2015 dataset, we estimated the proportion of infant at birth attributable to mother. For 4 month infants, we estimated the proportion attributable to the mother and itself at birth. For 12 month infants, we estimated the proportion attributable to the mother and itself at birth and four months (Shenhav et al. 2019). This allowed “unknown” to be more strictly defined as the component of the infant microbiome that could not be explained by the mother. It also allowed us to better discern if completely new strains were acquired at the 4^th^ and 12^th^ months of life (that were not already acquired during previous life stages).

First, consistent with previous findings made with species and SNVs (Nayfach et al., 2016), species-FEAST estimates an increasing contribution of the mother over time (t-test p-value = 5.1 × 10^-4^), but SNV-FEAST estimates a decrease over time (p-value = 0.063) (**Figure 3**).

**Figure 3.**
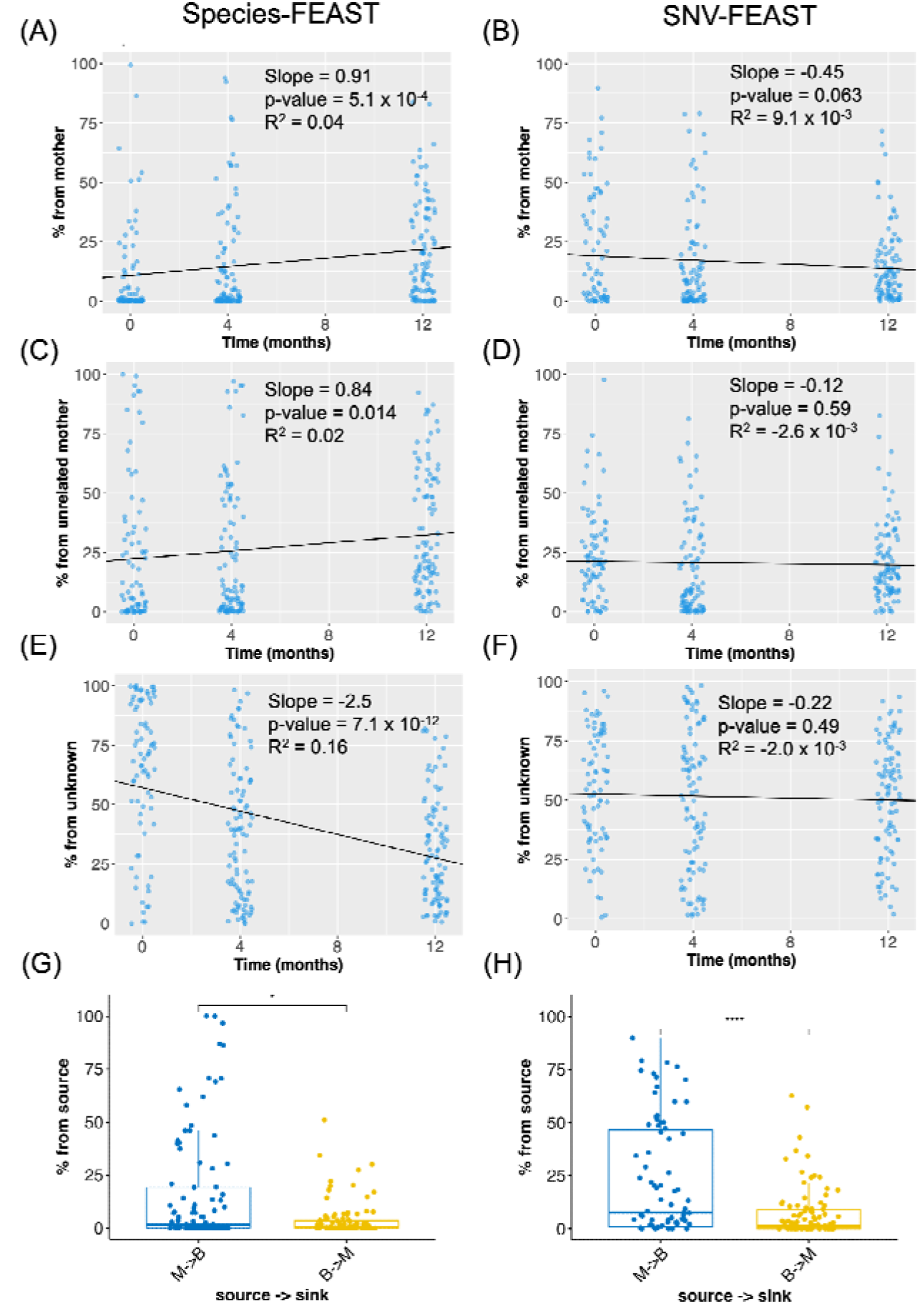
Source tracking in the infant gut microbiome over the first year of life. Species- and SNV-FEAST were applied to Backhed et al. 2019 data to estimate the contribution of (A, B) mother, (C, D) unrelated mothers and (E, F) unknown sources to infants sampled at birth, four months, and twelve months. The black line and inset statistics pertain to the linear regression fit for the source estimates as a function of age of the infant. (G, H) are flipped source tracking analyses with mother and infant swapped when using species-FEAST and SNV-FEAST, respectively. **Figure S4** shows the species that were included in species-FEAST and species that had SNVs included in SNV-FEAST. **Figure S5** shows the estimate of the unknown component when previous time points of the infant are excluded from the sources.

Second, we assessed the ability of species and SNV-FEAST to distinguish the true mother from three randomly selected unrelated mothers. Species-FEAST estimates an increasing contribution of unrelated mothers over time (t-test p-value = 0.014) while SNV-FEAST estimates no significant change over time (t-test p-value = 0.59) (**Figure 3**). The increase in contribution from unrelated mothers with species-FEAST does not suggest that these particular unrelated mothers are seeding the infant. Rather, the opposing trend observed with SNVs suggests that similarity at the species level is consistent with the maturation of the infant microbiome over time.

Finally, we estimated contributions from unknown sources, i.e. the proportion of the infant microbiome not explainable by the true mother, the three randomly selected unrelated mothers, or any previous time point. Species-FEAST estimates a sharp decline in contribution of unknown sources over the first year of life (t-test p-value =7.1 × 10^-12^) (**Figure 3**). This significant decrease in unknown at the species level reflects the infant microbiome maturation over the first year of life. By contrast, SNV-FEAST estimates little change in the contribution of unknown sources (t-test p-value = 0.49) (**Figure 3**). Note that this unknown component reflects what was gained since a previous time point. In other words, at 12 months, the infant on average acquired the same fraction of unknown as it did at 4 months and birth. When source tracking is run without including previous time points as sources, the unknown component increases over the first year of life for SNVs only (**Figure S5)**.

Next, we sought to understand the effect of swapping sink and source in the re-analysis of Backhed et al. 2015 data. In **Figure 3G and H**, the infant at birth is the potential source and mother is the sink. The estimated contribution from baby to mother is significantly smaller (species-FEAST: 11.9 difference, Wilcoxon rank sum test p-value = 0.013; SNV-FEAST: 16.0 difference, p-value = 2.2 × 10^-5^) compared to that of mother to baby. This trend may be suggestive, but is not conclusive, of directionality, whereby a less diverse source is seeded by a more diverse source.

### Contribution of the NICU built environment to infant microbiomes

Next, we re-analyzed a metagenomic dataset studying the contribution of the hospital environment to the infant gut microbiome in the neonatal intensive care unit (NICU) (Brooks et al. 2017). This dataset is composed of microbiomes of infant stool, as well as the NICU rooms of the same infants at frequently touched surfaces, sink basins, the floor, and isolette-top sampled over an 11-month period (Brooks et al., 2017). We applied SNV and species-FEAST to assess the contribution of the infant’s own NICU room as well as a different NICU room in the vicinity of the infant’s gut microbiome (**see Methods**).

Concordant with the findings of Brooks et al., both SNV and species-FEAST detected that the most common source contributing to the infant microbiome was the floor and isolette-top from the infant’s own room (**Figures 4A and B**). SNV-FEAST found Infant 18 also had large contributions from their own room’s touched surfaces at multiple time points (**Figure 4B**), which is consistent with a finding by Brooks et al. that three strains found in Infant 18 perfectly matched (> 99.999% average nucleotide identity) strains found in the touched surfaces samples of Infant 18’s own room. Lastly, both species-FEAST and SNV-FEAST found Infant 6’s microbiome was explained almost entirely by samples from a different room with SNV-FEAST finding a sizeable contribution from both the floor and isolette top and the sink basin in this different room. This is concordant with Brooks et al.’s finding of multiple cases of strain sharing across rooms of Infant 6 and 12 for the different surfaces. FEAST with both data types is able to quantify the extent to which Infant 6’s microbiome was influenced by strains present in the built environment.

**Figure 4:**
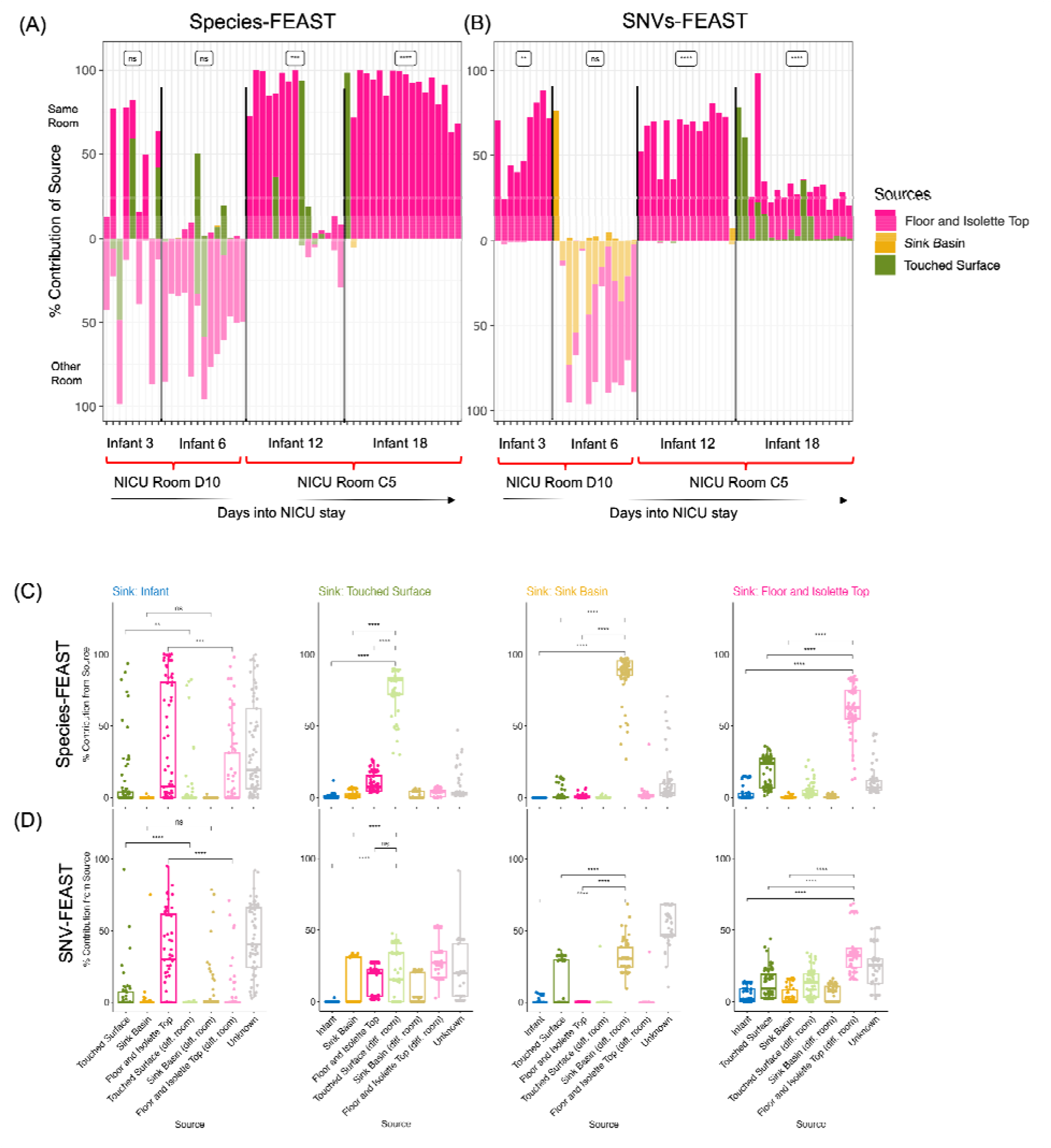
Source tracking of infant gut microbiome in the NICU. (A) species-FEAST and (B) SNV-FEAST applied to infants in the NICU. Each bar represents one sampling day in the NICU stay of an infant. Infants 3 and 6 stayed in the same room, but at different times. The same applies to Infants 12 and 18. The contribution of a different room was determined by using samples from Infant 12’s room for Infants 3 and 6, and samples from Infants 6’s room for Infants 12 and 18 for each of the categories of surfaces per infant: touched surface, sink basin, or floor and isolette top surface. The asterisks represent the result of a paired Wilcoxon signed rank test indicating whether the total contribution of surfaces from the infant’s own room were higher than contributions from the other room: **** for p-value < 0.0001, *** for p-value < 0.001, ** for p-value < 0.001, * for p-value < 0.05, and n.s. for p-value > 0.05. Iterative swapping of the infant sink and each potential source for source tracking with (C) species-FEAST and (D) SNV-FEAST. The first column shows source tracking results in which the infant was treated as the sink. In each column after the first column, a different environmental source was swapped with the infant and considered as a sink.

Through application of SNV and species-FEAST, we are able to quantify any trends over time the influence of the built environment on the infant microbiome (**Figures 4A and B**). SNV-FEAST more consistently finds that contribution from the infant’s own room exceeds contributions from a different room over time (paired Wilcoxon signed rank test for same room > different room: Infant 3: p-value = 1.95 × 10^-9^, Infant 6: 1, Infant 12: 3.05 × 10^-5^, Infant 18: 3.81 × 10^-6^) as compared to species-FEAST (Infant 3: p-value = 0.41, Infant 6: 1, Infant 12: 5.8 × 10^-4^, Infant 18: 3.81 × 10^-6^). Interestingly, species-FEAST assigns one dominant source primarily, whereas SNV-FEAST more often finds a combination of sources for a given sample.

Additionally, both SNV and species-FEAST estimated a large unknown component for all four infants, with Infant 18 showing the largest mean unknown component across the NICU stay based on SNVs (**Figure S6**). This unknown component is important because it signifies the extent to which other sources such as the mother and diet impact infant gut colonization.

We then asked the question: is the infant more explained by the built environment rather than vice-versa, the built environment is more explained by the infant. We tested this by swapping the infant and each of the three built environment sources (Figure 4C and D). The estimated contribution of room to infant is significantly higher than the estimated contribution of infant to room, but this asymmetry is more pronounced with SNV-FEAST. SNV-FEAST showed significantly higher contribution of room to infant for two of the three surface types (floor and isolette top: Wilcoxon rank sum test p-value = 7.00x 10^-9^, touched surface: p-value = 0.0058, sink basin: p-value = 0.274) while species-FEAST found this to be true for one of the three surface types (floor and isolette top: Wilcoxon rank sum test p-value = 7.1×10^-5^, touched surface: p-value = 0.968, sink basin: p-value = 0.998). Interestingly, the built environments of different rooms highly resemble each other. This is especially apparent with species-FEAST, suggestive of similar ecological forces operating in similar built environments. By contrast, SNV-FEAST reveals a higher diversity of contributing sources of the built environment samples to other NICU built environments, once again highlighting the utility of performing source tracking with SNVs.

### Global source tracking of ocean microbiomes

The ocean microbiome is a complex community that displays biogeography at the species and functional levels (Nayfach et al., 2016; Sunagawa et al., 2015). To further understand global patterns of ocean microbiomes, we applied SNV and species-FEAST to the Tara Oceans microbiome dataset (Sunagawa et al., 2015). In the source tracking context, rather than defining sharing as evidence of a transmission event (which is more likely in mother-infant data), estimated source contributions at best explain the extent to which a given ocean sample resembles other ocean samples. On one extreme, an ocean sample might be entirely explainable by a single ocean’s samples, and at the other extreme, an ocean sample might be explainable by multiple oceans at the same time. Another alternative is for an ocean sample to not be explainable by any of the provided sources, resulting in a high unknown component and potentially suggesting high endemism. These source tracking estimates could be indicative of the extent to which oceans mix or may be reflective of similar niches.

Tara Oceans is composed of 182 whole metagenomic sequencing samples derived from 64 stations at multiple depths. Previous research indicates that temperature is one of the highest drivers of variability in microbial composition in the ocean (Ladau et al., 2013; Sunagawa et al., 2015). For this reason, we restricted the source tracking analysis to sinks and sources from the same temperature and depth range: above 20 degrees Celsius and within an average of 5 meters below the surface.

First, we performed source tracking between oceans using SNV and species-FEAST. We treated each station around the world as a sink and estimated the contribution of different oceans around the world to that sink (**Methods**). Unknown represents any portion of the microbiome in these sink samples that cannot be explained by any of the provided source samples. We found that species and SNV-FEAST estimate different amounts of sharing between oceans, where SNVs estimate a higher unknown on average, potentially indicative of endemism. The finding that SNV-FEAST estimates a higher unknown contribution on average is most evident in the North Pacific, North Atlantic, South Atlantic, and Mediterranean oceans (**Figure S7**). Additionally, in some oceans, SNVs identify contributions from oceans that species-FEAST does not detect (**Figure 5, Figure S7**). For example, in applying FEAST to Indian Ocean samples we find that there is measurable sharing of microbes with the Mediterranean Sea, but this is not detected with species (**Figure 5C**). Such differences were found in samples from other oceans as well (**Figure S7**).

**Figure 5.**
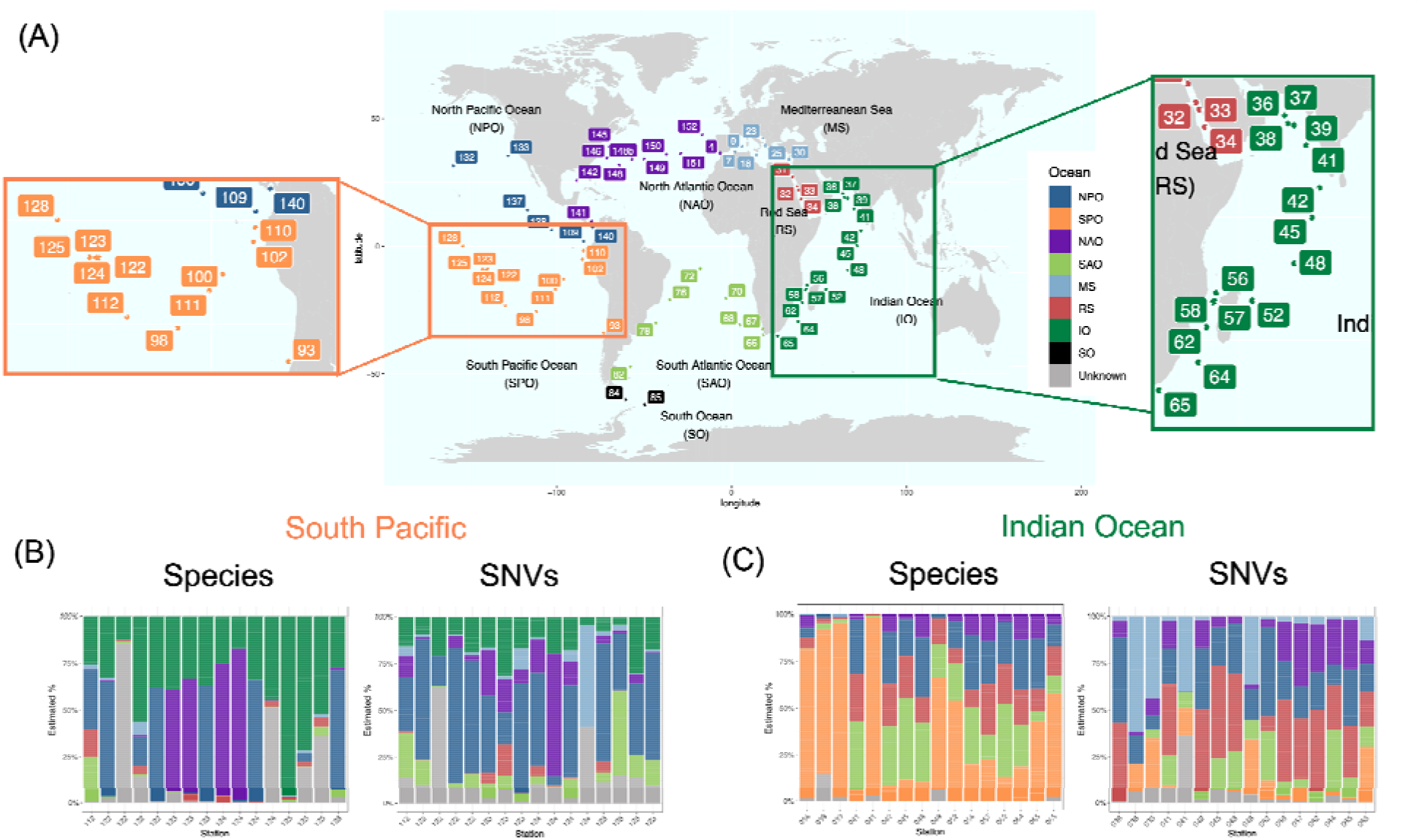
Microbial source tracking in the Tara Oceans dataset with SNV and species-FEAST. World map indicating the location of sampling sites (A). Source tracking estimates for the contribution of different oceans to the South Pacific (n=16) (B) and Indian Oceans (n=16) (C) are depicted with vertical bars. In each experiment, all stations around the world excluding those from the “sink” ocean are considered potential sources. Light blue, for example, represents the total contribution of the four stations from the Mediterranean Sea that had samples in the surface layer that were also greater than 20°C in temperature.

Next, we assessed whether source tracking estimates display a distance-decay relationship. Previous studies found that genetic distance, such as that represented by fixation index F_ST_, increases with geographic distance between populations (Cavalli-Sforza & Feldman, 2003; DeGiorgio & Rosenberg, 2013). Based on these findings, our expectation was that samples that are further away from a given station will have reduced resemblance to that station. To assess this distance-decay relationship, we plotted pairwiseh source tracking results across all possible pairs of ocean samples (**Figure 6A and B**). We found that indeed as the distance increases, the % explainability of a given source ocean decreases −0.23 % per thousand km according to species-FEAST (t-test p-value < 1 × 10^-16^), and −0.5% per thousand km according to SNV-FEAST (t-test p-value = 0.0018). The steeper slope for SNV-FEAST suggests that SNVs may be more sensitive to distance decay signals on a global level.

**Figure 6.**
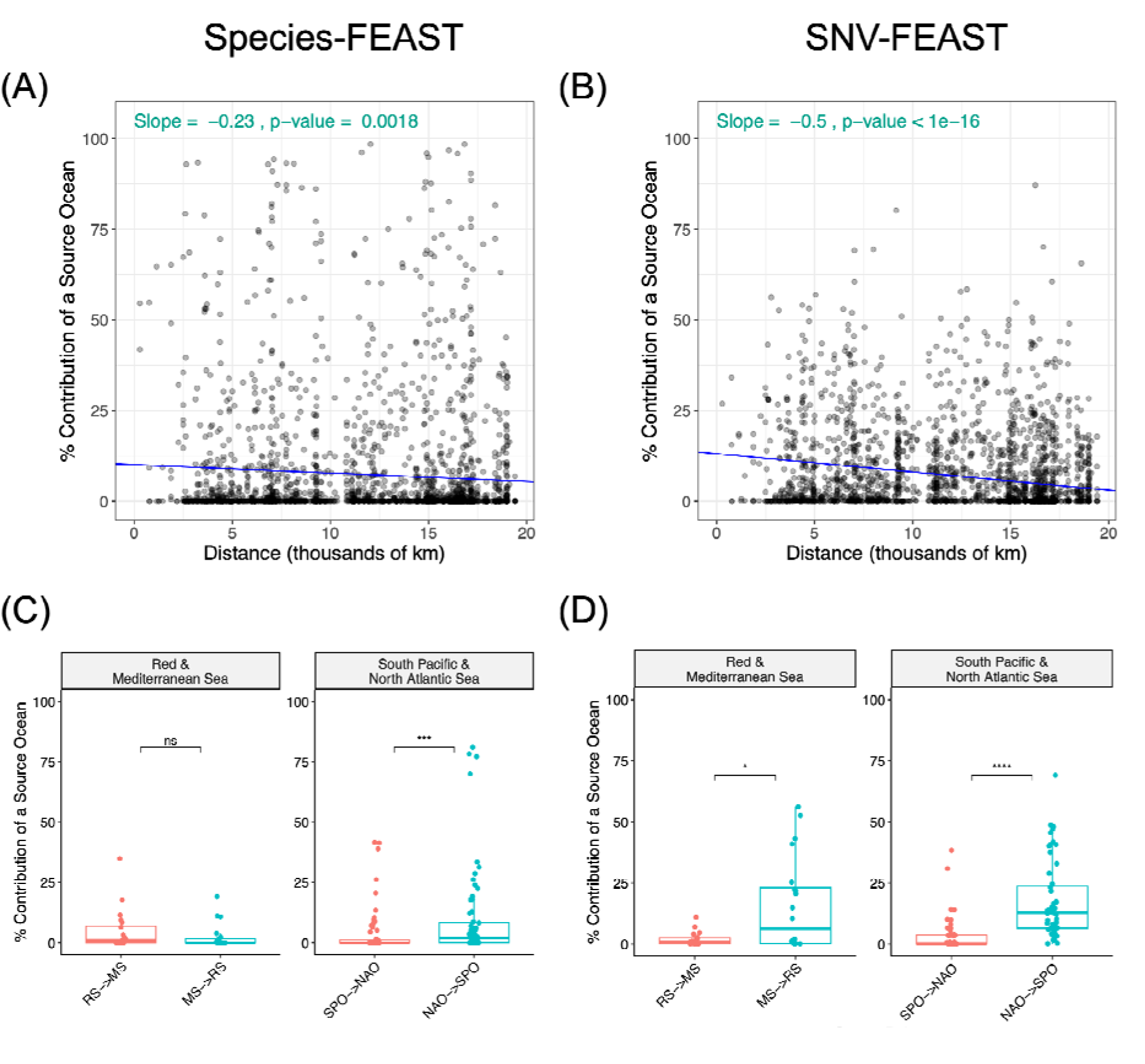
Source tracking with ocean samples. Distance decay in contribution of a “source” ocean to a “sink” ocean when using (A) species-FEAST and (B) SNV-FEAST. In each experiment, only stations from one ocean were considered as sources for a given sink station. For example, when performing source tracking between the mediterranean and north atlantic, for each mediterranean station, the 10 available north atlantic stations were considered as potential sources. Thus, plotted are 10 points for a given mediterranean sink, where each point represents the contribution of a source station from the North Atlantic to the Mediterranean sink station in question. Shown in inset text are the slope and t-test p-value for the slope. (C) and (D) are flipped source tracking analysis with the Red Sea and Mediterranean, as well as the South Pacific Ocean and North Atlantic Ocean using species-FEAST and SNV-FEAST, respectively.

Finally, we investigated whether some oceans have higher estimated contributions to other oceans than vice versa, potentially indicative of the directionality of transmissions (though see Discussion). Specifically, we investigated the relationship between the Red Sea to the Mediterranean Sea (**Figure 6C and D**). Migration from the Red Sea to the Mediterranean, known as Lessepsian migration, is well-documented for not only microorganisms but also macroorganisms like fish (Bentur et al., 2008; Bianchi & Morri, 2003; Golani, 2009). Anti-Lessepsian migration (Red Sea to Mediterranean), on the other hand, has been primarily thought to be rare due to the Additionally, recent studies suggest that there is also evidence for anti-Lessepsian migration of bacteria (Mediterranean to Red Sea) may be more common than Lessepsian migration (Elsaeed et al., 2021). Research studies find that Mediterranean has brine pools that produce similar a similar environment to the Red Sea’s (Antunes et al., 2011), allowing for bacteria from the MS to potentially thrive in the RS.

By swapping the Red Sea and Mediterranean as source and sink, we found that there was indeed a significant difference in the estimated contribution from one direction to another with SNVs but not species (**Figure 6C and D**). SNV-FEAST found the Mediterranean explained an average of 15% of the Red Sea, while the Red Sea explained an average of 1.8% of the Mediterranean (Wilcoxon rank sum test, p-value =0.02), consistent with anti-Lessepsian migration. Meanwhile, a similar analysis with species-FEAST found the Mediterranean explained 2.5% of the Red Sea and the Red Sea explained 4.9% of the Mediterranean (Wilcoxon rank sum test, p-value = 0.25). In a similar analysis between North Atlantic and South Pacific we found that both species and SNVs supported significantly greater contributions from the North Atlantic to the South Pacific, with SNV-FEAST estimating a greater contribution (17%, Wilcoxon rank sum test p-value = 5.1 × 10^-11^) than species-FEAST (10%, Wilcoxon rank sum test p-value =1.8 × 10^-4^).

Together, these results suggest that on average, SNV and species FEAST generate similar source tracking results in the Tara Oceans dataset, with SNVs displaying stronger signals of endemism, distance-decay relationships, and potential directionality of transmission.

## DISCUSSION

Source tracking provides insight into potential source contributions to a metagenomic sample as well as similarities between metagenomic samples. While species abundances have been informative in source tracking in several studies (Flores et al., 2011; Knights et al., 2011; McGhee et al., 2020; Shenhav et al., 2019), they may be limited in their resolution. SNVs provide a potential alternative because of their ability to distinguish sources of strain transmissions. Here we compared the ability of a previously published source tracking algorithm FEAST using species versus SNVs as input data. In application of species and SNV-FEAST to simulations as well as three case studies, we demonstrate that the two input types can provide distinct insights into microbial sharing and similarities across different environments. As a hypothetical example, two unrelated samples may have very similar species composition due to similar colonization processes and similar environmental influences without any actual microbial sharing. It would be unlikely for these two unrelated samples to share rare SNVs, however. This distinction suggests that SNVs indeed can provide insight into the ecological processes shaping microbial communities that species information alone cannot, and our three case studies are able to demonstrate this.

In the first case study, we confirmed previous findings that SNV sharing between mothers and infants decreases over the first year of life while species sharing increases (Nayfach et al., 2016), suggesting that while the infant microbiome matures to resemble adults at the species level, sources other than the mother may seed the infant over time. In the second case study, we confirmed source contributions from the NICU built environment to the infant microbiome (Brooks et al., 2017), and found that SNVs detect a more consistent estimate in source contributions overtime compared to species as well as detecting contribution from sources not detectable by species-FEAST.

In the third case study, we perform source tracking in the Tara oceans dataset and found SNVs display a stronger distance decay relationship. These distance-decay results parallel recent findings made with gene content (Dlugosch et al., 2022). While previous studies have examined the biogeography of the ocean using species profiles, genes (Dlugosch et al., 2022; Nayfach et al., 2016) or amino acid variants from a single species (SAR11) (Delmont et al., 2019), for the first time, we leverage the use of SNVs across all detected prevalent species in the ocean microbiome to identify proportions of sharing across oceans. A benefit of using SNVs in the ocean microbiome is that SNVs can track fragments of DNA that have moved due to horizontal gene transfer in the distant past rather than relying on inference of whole genomes or presence of private SNVs that may been transmitted in the recent past. This global-level source tracking is analogous to admixture estimation with human genotypes (Alexander et al., 2009; Chiu et al., 2022).

We note that source tracking provides insights into similarities between microbiomes and potential transmissions, though the directionality is less conclusive. It is possible that increased contributions in one direction but not the other is suggestive of directionality of transmission. For example, in the case of the mother-infant data from Backhed et al. 2015, FEAST predicted higher contribution from mother to baby than vice versa. This is consistent with work done on crAss-like phage transmissions between mother and infant in the same dataset that showed evidence of directionality by tracking the accumulation of mutations over time that are private to the infant and absent from the mother (Siranosian et al., 2020). But in the case of the ocean, it is possible that over longer time periods, differences in relative contributions from one part of the world to another (e.g. Mediterranean to Red Sea) are more reflective of local selection pressures that permit certain species and genotypes (Delmont et al., 2019). Thus, source tracking in certain instances, such as the ocean microbiome, at best reflects the extent of similarity between samples and is less conclusive about directionality.

A popular approach used to track strain transmissions is by detecting high average nucleotide identity (ANI) for species shared between source and sink. For example, inStrain (Olm et al., 2021) identifies a match between samples for a given species when ANI exceeds 99.999%. However, it is to be noted that inStrain provides distinct and complementary information from FEAST given its binarization of whether or not a strain is shared. For illustration purposes, if an infant harbors 100 species, of which only 1 came from their mother, but that species’ strain’s relative abundance is 50% of the infant’s microbiome, SNV-FEAST would infer that the mother’s contribution is 50%, while inStrain would infer that only 1/100^th^ of the species are derived from the mother.

Other studies rely on tracking transmissions of strains with private SNVs shared only between the sink and putative source (Bäckhed et al., 2015; Korpela et al., 2018; Nayfach et al., 2016; Schmidt et al., 2019). The private marker allele tracking approach in Nayfach et al. 2016 provides an improved estimate of true percentage of species that share some portion of their genome with putative sources compared to inStrain (**Figure S2, S3**). It is possible that requiring only 5% of marker alleles to be shared rather than a 99.999% ANI permits detection of horizontal gene transfers between lineages residing in mothers and infants (D. W. Chen & Garud, 2022; Vatanen et al., 2022). However, in FEAST, by using any SNV with an informative distribution across sources as determined by our signature scoring method, we are able to quantify the relative contribution of all the sampled environments and assign a proportion to these putative sources. Another advantage to FEAST is that the contribution of unknown sources can be quantified. For example, the significant fraction of marine biodiversity estimated to be ‘unknown’ may be endemic, as previously noted in the Mediterranean (Katsanevakis et al., 2014).

A drawback, however, with using SNVs over species is deeper, whole genome sequencing is required to accurately call SNVs. Moreover, even when there is sufficient coverage, there is still the challenge of a large number of SNVs. We demonstrate one way to subset SNVs that uses a scoring method for informativeness, but there may yet be other methods for filtering SNVs to the most informative set. Another potential caveat of SNV filtering is that not all species present will be represented in the final signature SNV set (**Figure S4**). Species with higher abundance are more likely to be represented in the signature SNV set. However, we show that not all species need to contribute signature SNVs in order to make accurate inferences, and likewise, not all SNVs are needed to make accurate inferences (**Figure S1**).

Ascertainment of SNVs from metagenomic data in a high-throughput manner, especially common SNVs with microbiome genotyping technology (Shi et al., 2021), is becoming an increasing priority for the field as metagenomic datasets become more abundant. A genotyper for prokaryotes has already been developed and tested on a catalog of over 100 million SNVs in order to characterize population structure (Shi et al., 2021). Such a catalog of informative SNVs could be invaluable for source tracking. With source tracking enabling us to characterize samples by their relationship to known samples, we have a powerful tool to explore samples in new contexts we have yet to discover.

## METHODS

### Data

For simulations and analyses of infant microbiomes in the first year of life (Bäckhed et al., 2015), we downloaded the raw shotgun metagenomic sequencing reads from public read archives under accession number PRJEB6456. We downloaded the raw sequence reads for the NICU analysis (Brooks et al., 2017) from accession number PRJEB323631, and the equivalent for the Tara Oceans analyses (Sunagawa et al., 2015) were downloaded from accession number PRJEB402. Data from the HMP Consortium (Methé et al., 2012) and Lloyd-Price et al (Lloyd-Price et al., 2017) was downloaded from the following URL: https://aws.amazon.com/datasets/human-microbiome-project/.

### Estimation of species and SNV content of metagenomic samples

We used MIDAS (Metagenomic Intra-Species Diversity Analysis System, version 1.2, downloaded on November 21, 2016 (Nayfach et al., 2016) to estimate species abundance and SNV content per species in each metagenomic shotgun sequencing sample. The database we used to apply MIDAS consisted of 31,007 bacterial genomes that are clustered into 5,952 species. The parameters we used to estimate species abundances and SNVs were described in (Garud et al., 2019). A species was considered present if there are at least 3 reads mapping to a set of single copy marker genes on average. To call SNVs, we used the default MIDAS settings in order to map reads to a single representative reference genome. The mapping was done with Bowtie 2 (Langmead & Salzberg, 2012): global alignment, MAPID≥94.0%, READQ≥20, ALN_COV>0.75, and MAPQ≥20, where species with reads mapped to less than 40% of the genome were excluded from the SNV calls. We excluded samples with depth lower than 5 reads, and excluded genetic sites using the default site filters of MIDAS (e.g. ALLELE_FREQ>0.01, with the exception of SITE_DEPTH which was set to 3.

### Application of FEAST algorithm

FEAST, originally introduced by Shenhav et al., is an R-based method that models the mixture proportions for various “source” microbial samples for a given “sink” (Shenhav et al., 2019). This method utilizes expectation maximization to estimate the proportions when given any sort of count-based feature matrix representing the potential sources and sinks. The intuition behind the estimation process is that a source with a similar species distribution to the sink would have a higher contribution estimate to the sink. A species with non-zero counts only in source *j* and the sink would increase the estimated contribution of source *j*. However, in many cases, the same species are found in multiple sources simultaneous. The algorithm does not uniquely assign a species to a source but rather simultaneously utilizes all species information to infer the source contributions. The method was originally tested and evaluated on species and not on more fine scale genetic data such as SNVs. The number of different species, on average, range in number from a few hundred to a few thousand, while the number of possible nucleotide sites that vary across different sources can number in millions. For this reason, a SNV-filtering process is necessary so that the algorithm can run within a reasonable time and with reasonable memory requirements.

### Application of FEAST to the Backhed et al. 2015 dataset

For both species and SNV-FEAST, the same set of sources and sinks were fed into the FEAST algorithm. In the case study of infants in the first year of life (Bäckhed et al., 2015), the sink consisted of the infant fecal sample at either four days, four months, or 12 months and the potential sources consisted of fecal samples from the true mother, three randomly selected mothers from the same dataset, and also any previous time points for the infant.

Species-FEAST utilized all species present in the infant whereas SNV-FEAST used signature SNVs from the subset of species that had signature SNVs. Shown in **Figure S3** are the distribution of species included in species and SNV-FEAST.

### Application of FEAST to the Brooks et al. 2017 dataset

For the case study of infants in the NICU (Brooks et al., 2017), the sink consisted of the fecal sample of the infant at a given time point and the potential sources consisted of pooled reads from the touched surfaces, the sink basin and the floor and isolette top from both the infant’s own room as well as a different room. The different room was Infant 12’s room for Infants 3 and 6, Infants 6’s room for Infants 12 and 18.

### Application of FEAST to the Sunagawa et al. 2015 dataset: Determining the signature SNV set

Signature SNVs were identified as described in the main text. We provide specific steps for determining signature SNVs:

1. Filter sites: only sites of the genome with at least the required number of reads mapping to the site are considered. In the case study of infants in the first year of life (Bäckhed et al., 2015) and infants in the NICU (Brooks et al., 2017), the minimum coverage requirement is 10 across the sink and *J* sources. For the Tara Ocean (Sunagawa et al., 2015) samples, the minimum coverage is five reads (Sunagawa et al., 2015). Additionally, sites that are biallelic must have more than one read mapped to each allele to be considered.
2. Perform per site per source parameter estimates: for each potential source compute the estimated allele frequency in the sink θ under two different hypotheses: **Hypothesis 1:** Source *i* with allele frequency *p_i_* explains the allele counts in the sink.

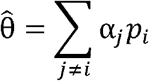 **Hypothesis 2:** A combination of all other sources except *i* (sources *j* ≠ *i*) explain the observed allele count distribution in the sink. The estimate of the sink allele frequency is computed using a mixture of the allele frequencies *p_j_* from those sources. The mixing parameter *α_j_* is learned using Sequential Least Squares Programming (scipy.minimize()) with the constraint of summing to 1 with bounds of 0 to 1 inclusive: ∑_*j*≠*i*_ *α_j_* = 1.
3. Compute per site per source log likelihoods: Compute the binomial log-likelihood under hypotheses 1 and 2, given *n* reads with the reference allele and *m* reads with the alternative allele in the sink:

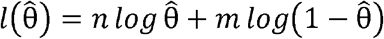
4. Compute per site per source log likelihood ratio:

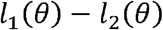
5. Compute per site summary signature score: The maximum log likelihood ratio per site is the signature score for that SNV, representing how favorably one of the sources explains the sink over all other sources
6. Filtering of SNVs using signature score: One signature score for that SNV represents how favorably one source explains the sink better than all other sources. All the scores are ranked across SNVs and SNVs with scores that are greater than two standard deviations over the mean signature score within each 200 kbp window of the genome are retained as signature SNVs. This window size was chosen for to optimize run time and memory requirements.

Note, if only one source passes minimum coverage filtering, *l*_2_ (*θ*) – 0 resulting in a very high likelihood ratio as represented by *l*_1_(*θ*) for the one source. These SNVs are more likely to pass the signature score filtering. One exception for SNVs that are included in the signature SNV set without passing signature score filtering are SNVs with an allele that is completely unique to the infant, as these represent SNVs that are potentially derived from an unknown source. Signature SNVs are obtained from the SNV profile of every species for which there is MIDAS output.

### Simulating mother to infant transmission

The mixture proportions for 28 simulated infants is shown in **Table S1**. Four possible scenarios are simulated using a combination of either low or high number of sources and low or high transmission probabilities of species. High transmission of species was simulated by drawing separate transmission probabilities for each species in each contributing source based on a beta distribution with a mean equal to the species relative abundance and variance equal to 0.1, a value selected to emulate Backhed et al.’s mean relative abundance and variance. For the low transmission scenario, transmission probabilities were drawn from a beta distribution with mean 0.1 times the relative abundance and variance at 0.1. To determine if a species from each source was transmitted to a given infant, a binomial draw was performed *J* times, where *J* = number of sources, and the probability of a mother transmitting the species is *p_j_* based on the beta-drawn transmission probability. If any of the draws yields a one, that species is transmitted to the infant from all sources. The same simulated data under these scenarios is used for both SNV and species source tracking.

The source tracking estimates are compared to the true mixing proportions using Spearman correlation. The significance of correlation is calculated using the stat_cor function in the ‘ggpubr’ package (*CRAN - Package Ggpubr*, n.d.).

### Comparison to inStrain

We ran inStrain (Olm et al., 2021) on the same synthetic samples as described above. InStrain “profile” (Olm et al., 2021) and inStrain “compare” (Olm et al., 2021) were run for every possible infant-source pair. For example, for simulated infant 1 there were 10 putative sources, therefore inStrain compare was run 10 times for each putative source. InStrain reports popANI calculated per scaffold for a given species. To compute a single statistic per species, we computed the average popANI across scaffolds for a given species. The percent infant microbiome species that had strains shared with mother was computed as the number of species in which popANI was >= 99.999% divided by the total number of species with coverage >= 5. PopANI was only calculated in scaffolds that had >=5 coverage in both samples of the pair.

### Comparison with strain tracking approach in Nayfach et al. 2016

We applied the strain tracking approach in Nayfach et al. 2016 on the same synthetic communities described above. In Nayfach et al. 2016, strain transmissions are tracked by identifying ‘marker alleles’ which are private to the infant, mother, or infant-mother dyad, and absent from the broader population. A strain is considered to be shared if at least 5% of all marker alleles for a mother-infant dyad are shared. Note that the approach for strain tracking proposed in Nayfach et al. 2016 utilizes SNV information outputted by MIDAS, but is not a part of MIDAS.

Each simulated infant had up to 10 sources that were real maternal samples from Backhed et al. 2015 For each possible pair of infants and maternal sources (10 pairings per infant, with 48 infants), we found the set of infant-only marker alleles, mother-only marker alleles, and mother-infant dyad marker aleles. As described in Nayfach et al, 2016, only sites with minimum 30 reads and only alleles that were supported by at least 10% of the total reads aligned to that site were considered. The infant marker allele and mother marker allele were defined as alleles that were present only in the focal sample and absent from the background samples (or below 3 reads = 10% * 30 reads). For the infant, the background consisted of all mothers (including mothers that were used to simulate the infant), real infant samples (excluding infants of mothers used to simulate the infant), and 337 samples of adults from the United States in the HMP (which includes 180 unique adults) that were obtained from the metagenomics repository of HMP under project ID SRP002163 and SRP056641 (Lloyd-Price et al., 2017; Methé et al., 2012). For the mother, the background consisted of all mother and infant samples in addition to the HMP samples. For computing shared marker alleles, an allele must be present in both the mother and infant but absent from the background, which consisted of all mothers and the HMP samples.

To compute sharing, two quantities were considered: “total sharing”, defined as % shared marker alleles/ (infant marker alleles + mother marker alleles + shared marker alleles) and proportion of infant marker alleles that are shared: % shared marker alleles/ (infant marker alleles + shared marker alleles). The first quantity compared to FEAST estimates was the percentage of infant species in which the “total sharing” was at least 5%. The second quantity compared to FEAST was the pooled proportion of infant marker alleles that are shared across all species.

### Distance Decay Analysis

To study the relationship between source tracking estimates and geographic distance, we analyzed all oceans as either a sink or source against all other possible oceans. To compute geographic distance between stations, we applied the Haversine distance to the longitude and latitude of the sampling sites provided by (Sunagawa et al., 2015) using the package “geosphere” (Hijmans et al., 2021). Source tracking estimates were computed as described above using either SNV-FEAST or Species FEAST. The regression line for the distance decay analysis was computed using a linear mixed model “contribution ~ distance + (11 sink_ocean)”.

## Ethics declarations

The authors declare that they have no competing interests.

## Acknowledgements

We thank Nicole Zeltser for processing the Tara Oceans data through the MIDAS pipeline. We thank Richard Wolff for his advice on simulations. We thank Michael Wasney for testing the associated software. We thank members of the Garud lab for their feedback on the manuscript. LB was supported by the NSF Graduate Research Fellowship Program grant numbers DGE-1650604 and DGE-2034835. N.R.G. received support from the Paul Allen Frontiers Group and the Research Corporation for Science Advancement.

## Availability of Data and Materials

All metagenomic data was obtained from public repositories. The applicable accessions numbers are PRJEB6456 for Backhed et al. 2015 (mother-infant), PRJEB323631 for Brooks et al. 2017 (NICU), PRJEB402 for Sunagawa et al. 2015 (Tara Oceans), and SRP002163 and SRP056641 for HMP.

Source code for SNV-FEAST signature SNV selection as well as analyses in this paper are available at GitHub (https://github.com/garudlab/Signature-SNVs), Zenodo (DOI 10.5281/zenodo.7515044), and PyPi for pip installation (https://pypi.org/project/Signature-SNVs/0.0.1/).

## Notes

https://github.com/garudlab/Signature-SNVs

### Competing Interest Statement

The authors have declared no competing interest.

### Summary of Updates

We clarify the meaning and caveats of source tracking throughout the manuscript. We have added extensive new simulations in which we explore the abilities of SNV and species-FEAST and compare their abilities to strain-tracking approaches. Additionally, we added experiments that demonstrate that the contributions from one sample to another (e.g. mother to infant) are asymmetric. We have also substantially revised our ocean analysis and demonstrate significant distance decay trends in the ocean. A package for our Signature SNV approach is now made available through links in the manuscript.

