## Supplemental Information for "SNV-FEAST: microbial source tracking with single nucleotide variants"

**Contents:**

**List of Tables:**

| Table 1 | Mixture proportions for simulated gut microbiome of infants | Page 2 |
| --- | --- | --- |

**List of Figures:**

| Fig.S1  Fig.S2  Fig.S3 | Robustness testing of SNV-FEAST  Comparison of SNV-FEAST with inStrain using simulations  Comparison of SNV-FEAST with Nayfach using simulations | Page 3  Page 4  Page 5 |
| --- | --- | --- |
| Fig.S4 | Species with signature SNVs in Backhed et al. data | Page 6 |
| Fig.S5  Fig.S6  Fig.S7  Fig.S8 | Microbial source tracking with infants in the first year of life.  Microbial source tracking with infants in the NICU and their built environment.  Microbial source tracking in the Tara Oceans dataset with SNV and species-FEAST.  Flipped source tracking for all ocean pairs | Page 7  Page 8  Page 9  Page 10 |

|  |  | **Source 1** | **Source 2** | **Source 3** | **Source 4** | **Source 5** | **Source 6** | **Source 7** | **Source 8** | **Source 9** | **Source 10** | **Unknown** |  | **Unknown** |
| --- | --- | --- | --- | --- | --- | --- | --- | --- | --- | --- | --- | --- | --- | --- |
| Complex Sink | **Trial 1** | 0.2 | 0.15 | 0.05 | 0.05 | 0.05 | 0.05 | 0.05 | 0.05 | 0.05 | 0.05 | 0.25 |  | 0-30% |
|  | **Trial 2** | 0.3 | 0.1 | 0.08 | 0.08 | 0.08 | 0.06 | 0.05 | 0.05 | 0.05 | 0.05 | 0.1 |  | 30-70% |
|  | **Trial 3** | 0.16 | 0.12 | 0.07 | 0.05 | 0.04 | 0.04 | 0.03 | 0.03 | 0.02 | 0 | 0.44 |  | 70-90% |
|  | **Trial 4** | 0.21 | 0.18 | 0.11 | 0.1 | 0.1 | 0.05 | 0.05 | 0.05 | 0.05 | 0 | 0.1 |  |  |
|  | **Trial 5** | 0.23 | 0.07 | 0.05 | 0.05 | 0.05 | 0.05 | 0.05 | 0.05 | 0.05 | 0 | 0.35 |  |  |
|  | **Trial 6** | 0.17 | 0.13 | 0.05 | 0.05 | 0.05 | 0.05 | 0.05 | 0.05 | 0 | 0 | 0.4 |  |  |
|  | **Trial 7** | 0.24 | 0.16 | 0.13 | 0.1 | 0.05 | 0.05 | 0.05 | 0.05 | 0 | 0 | 0.17 |  |  |
|  | **Trial 8** | 0.2 | 0.17 | 0.06 | 0.06 | 0.06 | 0.05 | 0.05 | 0 | 0 | 0 | 0.35 |  |  |
|  | **Trial 9** | 0.4 | 0.16 | 0.06 | 0.06 | 0.06 | 0.06 | 0.05 | 0 | 0 | 0 | 0.15 |  |  |
|  | **Trial 10** | 0.09 | 0.03 | 0.03 | 0.02 | 0.02 | 0.02 | 0 | 0 | 0 | 0 | 0.79 |  |  |
|  | **Trial 11** | 0.17 | 0.15 | 0.12 | 0.1 | 0.1 | 0.05 | 0 | 0 | 0 | 0 | 0.31 |  |  |
|  | **Trial 12** | 0.17 | 0.15 | 0.12 | 0.1 | 0.1 | 0.05 | 0 | 0 | 0 | 0 | 0.31 |  |  |
|  | **Trial 13** | 0.2 | 0.17 | 0.1 | 0.1 | 0.1 | 0.05 | 0 | 0 | 0 | 0 | 0.28 |  |  |
|  | **Trial 14** | 0.25 | 0.15 | 0.11 | 0.06 | 0.05 | 0.05 | 0 | 0 | 0 | 0 | 0.33 |  |  |
|  | **Trial 15** | 0.37 | 0.12 | 0.1 | 0.09 | 0.09 | 0.07 | 0 | 0 | 0 | 0 | 0.16 |  |  |
|  | **Trial 16** | 0.42 | 0.11 | 0.1 | 0.09 | 0.09 | 0.07 | 0 | 0 | 0 | 0 | 0.12 |  |  |
|  | **Trial 17** | 0.42 | 0.11 | 0.1 | 0.07 | 0.06 | 0.06 | 0 | 0 | 0 | 0 | 0.18 |  |  |
|  | **Trial 18** | 0.45 | 0.1 | 0.1 | 0.09 | 0.09 | 0.07 | 0 | 0 | 0 | 0 | 0.1 |  |  |
|  | **Trial 19** | 0.22 | 0.2 | 0.11 | 0.08 | 0.05 | 0 | 0 | 0 | 0 | 0 | 0.34 |  |  |
|  | **Trial 20** | 0.23 | 0.05 | 0.04 | 0.03 | 0.02 | 0 | 0 | 0 | 0 | 0 | 0.63 |  |  |
|  | **Trial 21** | 0.29 | 0.1 | 0.04 | 0.04 | 0.03 | 0 | 0 | 0 | 0 | 0 | 0.5 |  |  |
|  | **Trial 22** | 0.37 | 0.03 | 0.03 | 0.03 | 0.03 | 0 | 0 | 0 | 0 | 0 | 0.51 |  |  |
|  | **Trial 23** | 0.41 | 0.26 | 0.1 | 0.05 | 0.05 | 0 | 0 | 0 | 0 | 0 | 0.13 |  |  |
| Simple Sink | **Trial 24** | 0.2 | 0.18 | 0.09 | 0.09 | 0 | 0 | 0 | 0 | 0 | 0 | 0.44 |  |  |
|  | **Trial 25** | 0.37 | 0.26 | 0.21 | 0.05 | 0 | 0 | 0 | 0 | 0 | 0 | 0.11 |  |  |
|  | **Trial 26** | 0.19 | 0.06 | 0.04 | 0 | 0 | 0 | 0 | 0 | 0 | 0 | 0.71 |  |  |
|  | **Trial 27** | 0.32 | 0.2 | 0.07 | 0 | 0 | 0 | 0 | 0 | 0 | 0 | 0.41 |  |  |
|  | **Trial 28** | 0.55 | 0.14 | 0.08 | 0 | 0 | 0 | 0 | 0 | 0 | 0 | 0.23 |  |  |
|  | **Trial 29** | 0.75 | 0.15 | 0.05 | 0 | 0 | 0 | 0 | 0 | 0 | 0 | 0.05 |  |  |
|  | **Trial 30** | 0.85 | 0.05 | 0.05 | 0 | 0 | 0 | 0 | 0 | 0 | 0 | 0.05 |  |  |
|  | **Trial 31** | 0.06 | 0.04 | 0 | 0 | 0 | 0 | 0 | 0 | 0 | 0 | 0.9 |  |  |
|  | **Trial 32** | 0.13 | 0.1 | 0 | 0 | 0 | 0 | 0 | 0 | 0 | 0 | 0.77 |  |  |
|  | **Trial 33** | 0.16 | 0.04 | 0 | 0 | 0 | 0 | 0 | 0 | 0 | 0 | 0.8 |  |  |
|  | **Trial 34** | 0.36 | 0.23 | 0 | 0 | 0 | 0 | 0 | 0 | 0 | 0 | 0.41 |  |  |
|  | **Trial 35** | 0.72 | 0.14 | 0 | 0 | 0 | 0 | 0 | 0 | 0 | 0 | 0.14 |  |  |
|  | **Trial 36** | 0.1 | 0 | 0 | 0 | 0 | 0 | 0 | 0 | 0 | 0 | 0.9 |  |  |
|  | **Trial 37** | 0.17 | 0 | 0 | 0 | 0 | 0 | 0 | 0 | 0 | 0 | 0.83 |  |  |
|  | **Trial 38** | 0.31 | 0 | 0 | 0 | 0 | 0 | 0 | 0 | 0 | 0 | 0.69 |  |  |
|  | **Trial 39** | 0.4 | 0 | 0 | 0 | 0 | 0 | 0 | 0 | 0 | 0 | 0.6 |  |  |
|  | **Trial 40** | 0.48 | 0 | 0 | 0 | 0 | 0 | 0 | 0 | 0 | 0 | 0.52 |  |  |
|  | **Trial 41** | 0.5 | 0 | 0 | 0 | 0 | 0 | 0 | 0 | 0 | 0 | 0.5 |  |  |
|  | **Trial 42** | 0.61 | 0 | 0 | 0 | 0 | 0 | 0 | 0 | 0 | 0 | 0.39 |  |  |

**S1 Table. Mixing proportions for simulated infants.** To simulate complex (N sources > 5) and simple (N sources <= 5) sinks, we mixed varying proportions of reads from the FASTA files of real adult mothers extracted from the Backhed et al. 2015 dataset. Proportions shown represent the proportion of 10 million reads in infants that are taken from each source.

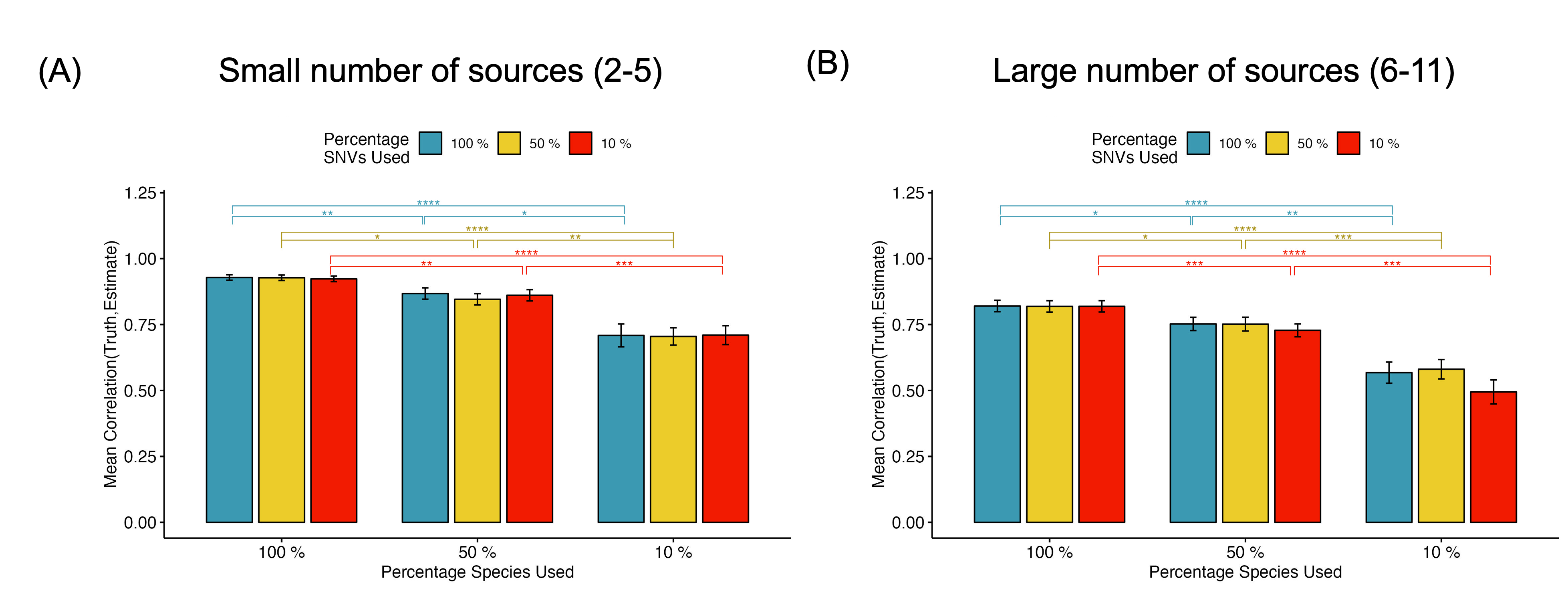

**Figure S1: Performance of SNV-FEAST as a function of fraction of species and SNVs included for analysis.** To assess whether all species and all signatures SNVs in the sink are needed for accurate source tracking with SNV-FEAST, we varied the proportion of species (from 10%, 50% or 100%) and SNVs (from 10%, 50% or 100%) included as inputs to the algorithm. The Y-axis values are Pearson Correlations between the estimated and true source tracking proportions. The errors bars represent the standard error of the mean. (A) Simulations with small number of contribution sources. (B) Simulations with a large number of contributing sources.

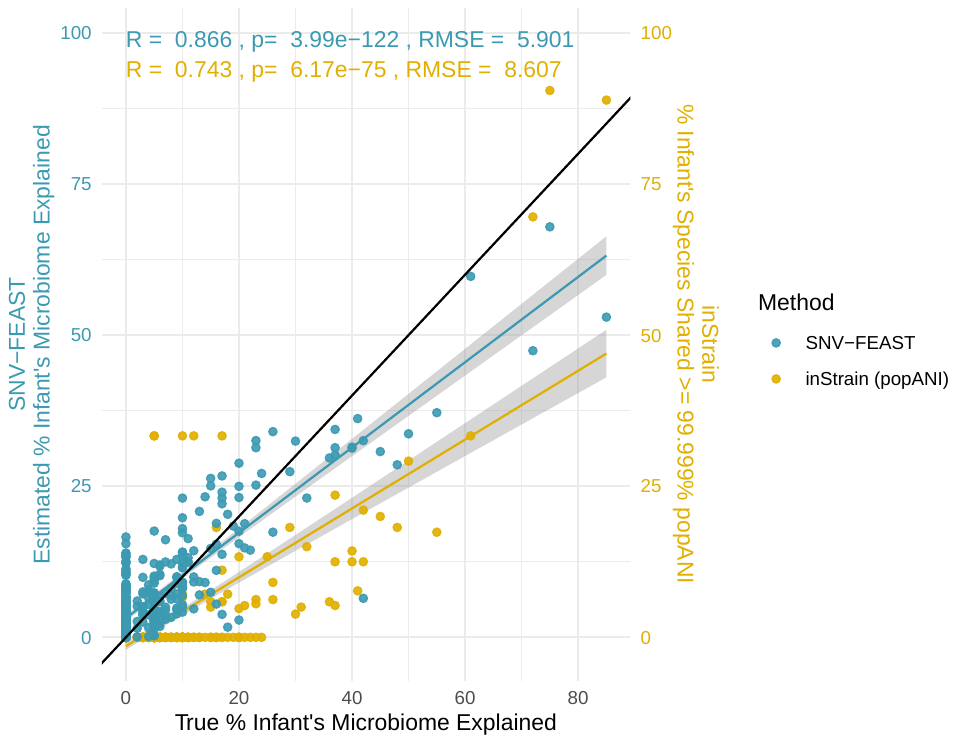

**Figure S2: Comparison of SNV-FEAST with inStrain.**

Application of SNV-FEAST and inStrain on simulated infant gut microbiomes in which the number of contributing sources was varied from 2 to 11 and the percentage of those contributing sources was varied from 1% to 90%. The x-axis represents the true proportion of the infant seeded by the source. Each point represents an infant-source pair. In the case of SNV-FEAST, the y-value represents the source tracking estimate. In the case of inStrain, the y-value represents the fraction of species in the infant that have at least 99.999% popANI with the source. Shown in the inset text is Pearson correlation and corresponding p-value and RMSE for both approaches.

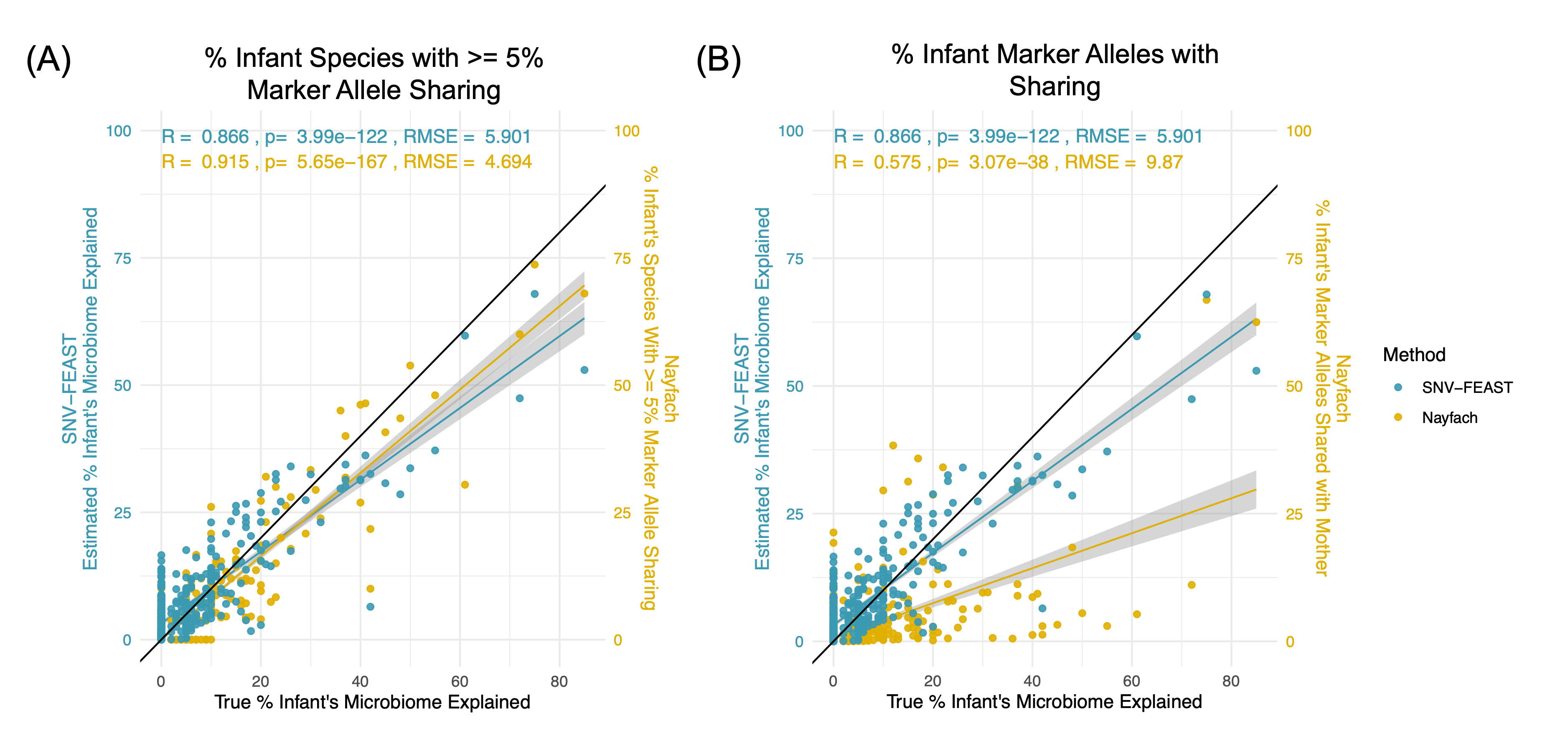

**Figure S3: Comparison of SNV-FEAST with the strain tracking approach in Nayfach et al. 2015.**

Application of SNV-FEAST and Nayfach et al. 2016 on simulated infant gut microbiomes in which the number of contributing sources was varied from 2 to 11 and the percentage of those contributing sources was varied from 1% to 90%. The x-axis represents the true proportion of the infant seeded by the source. Each point represents an infant-source pair. In the case of SNV-FEAST, the y-value represents the source tracking estimate. In the case of Nayfach et al. 2016, the y-value in (A) represents the fraction of species in the infant have at least 5% marker allele sharing while the y-value in (B) represents the fraction of all marker alleles in the infant that are shared with a given mother. Shown in the inset text is Pearson correlation and corresponding p-value and RMSE for both approaches.

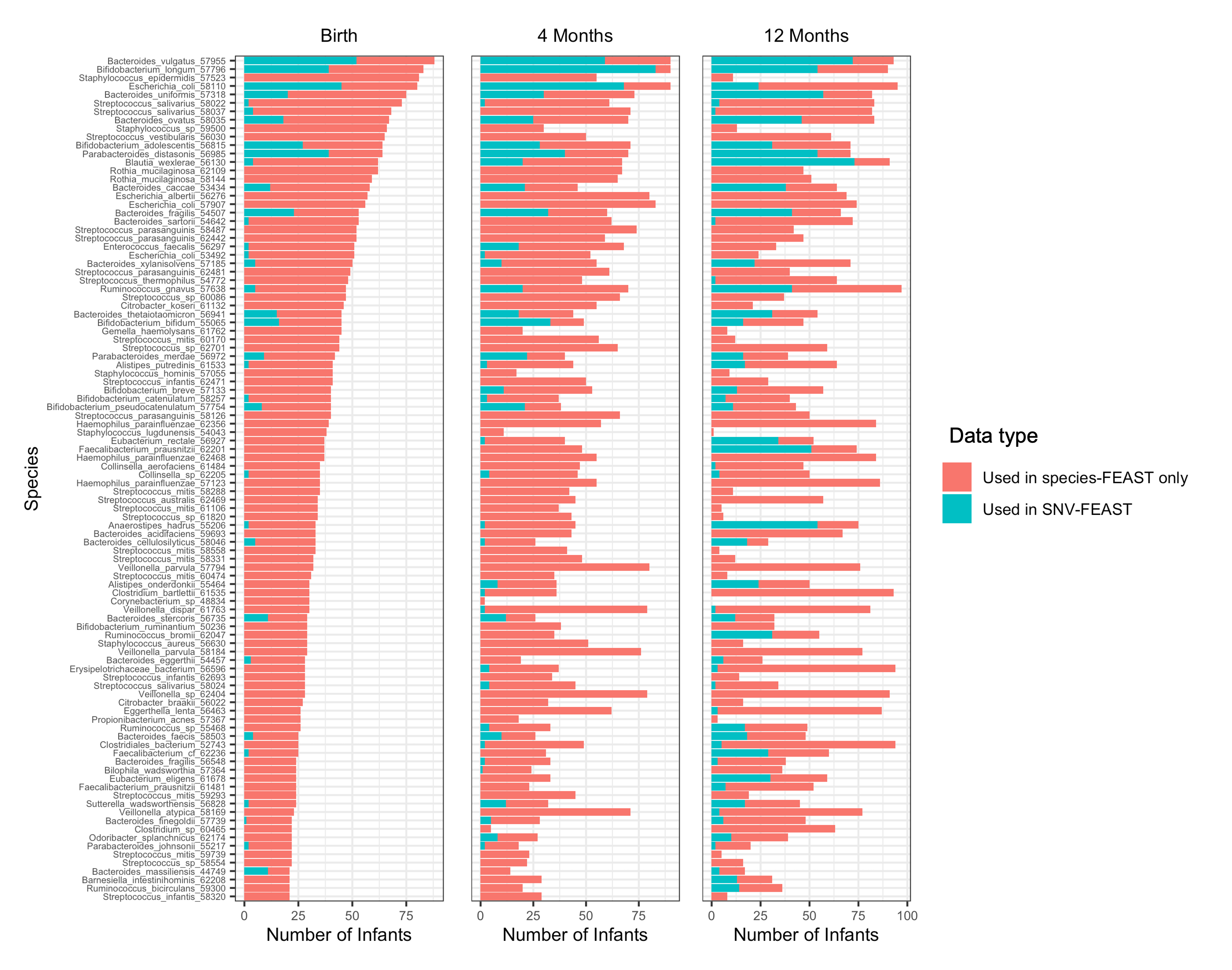

**Figure S4: Species with signature SNVs.** Number of infants in which certain species are detected in microbiome samples (whole bar) and in the signature SNV set obtained from those samples (teal bar) while the remained represents infants in which the species was only utilized in species-FEAST (salmon bar). Displayed are the 100 most prevalent species based on samples obtained from infants at birth.

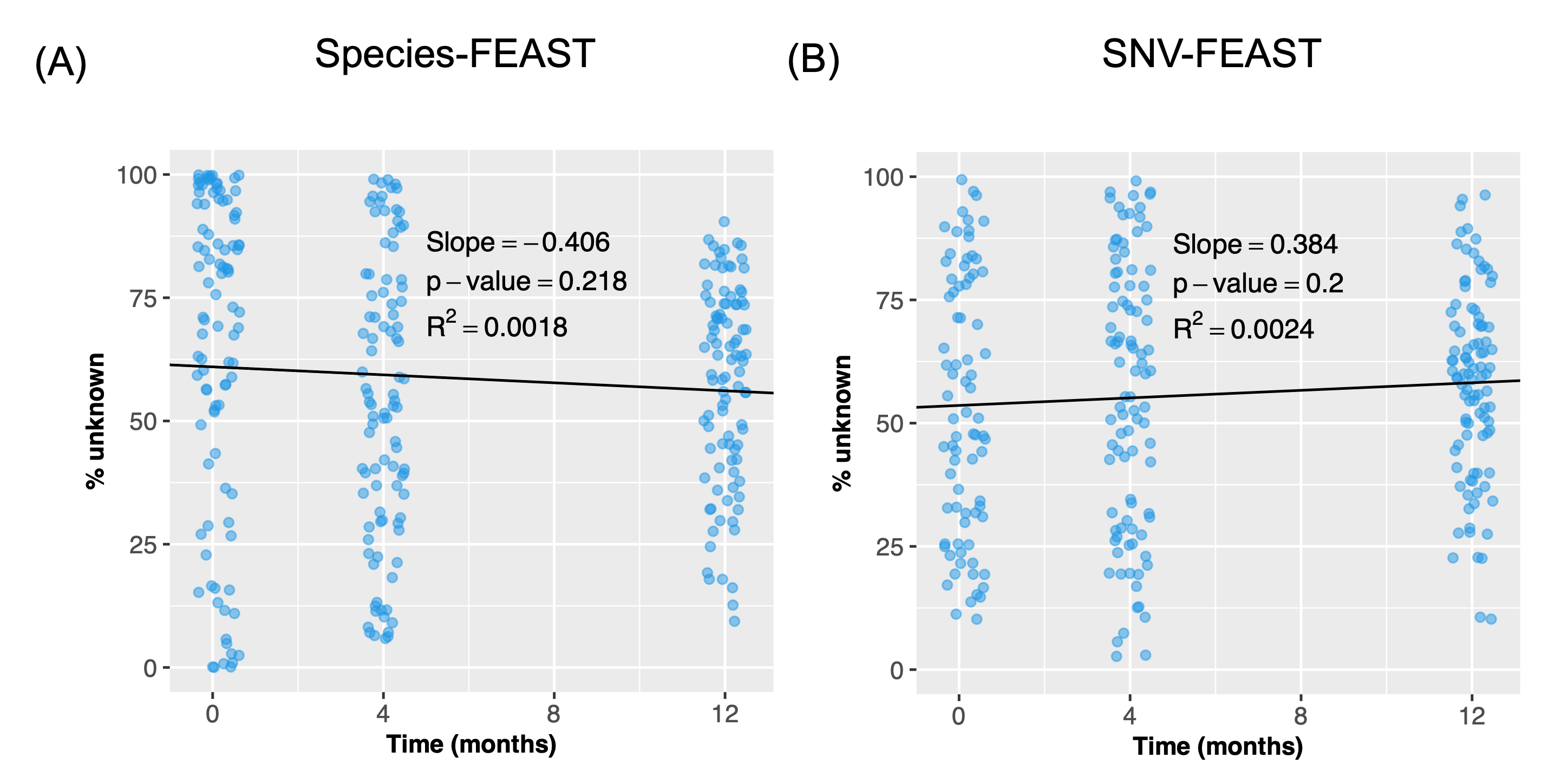

**Figure S5: Unknown component in microbial source tracking with infants in the first year of life.** Contribution of only unknown sources to the infant’s gut microbiome at birth, four and 12 months when previous time points of the infant are excluded as sources. Note this is a different experiment from the one shown in Figure 3.

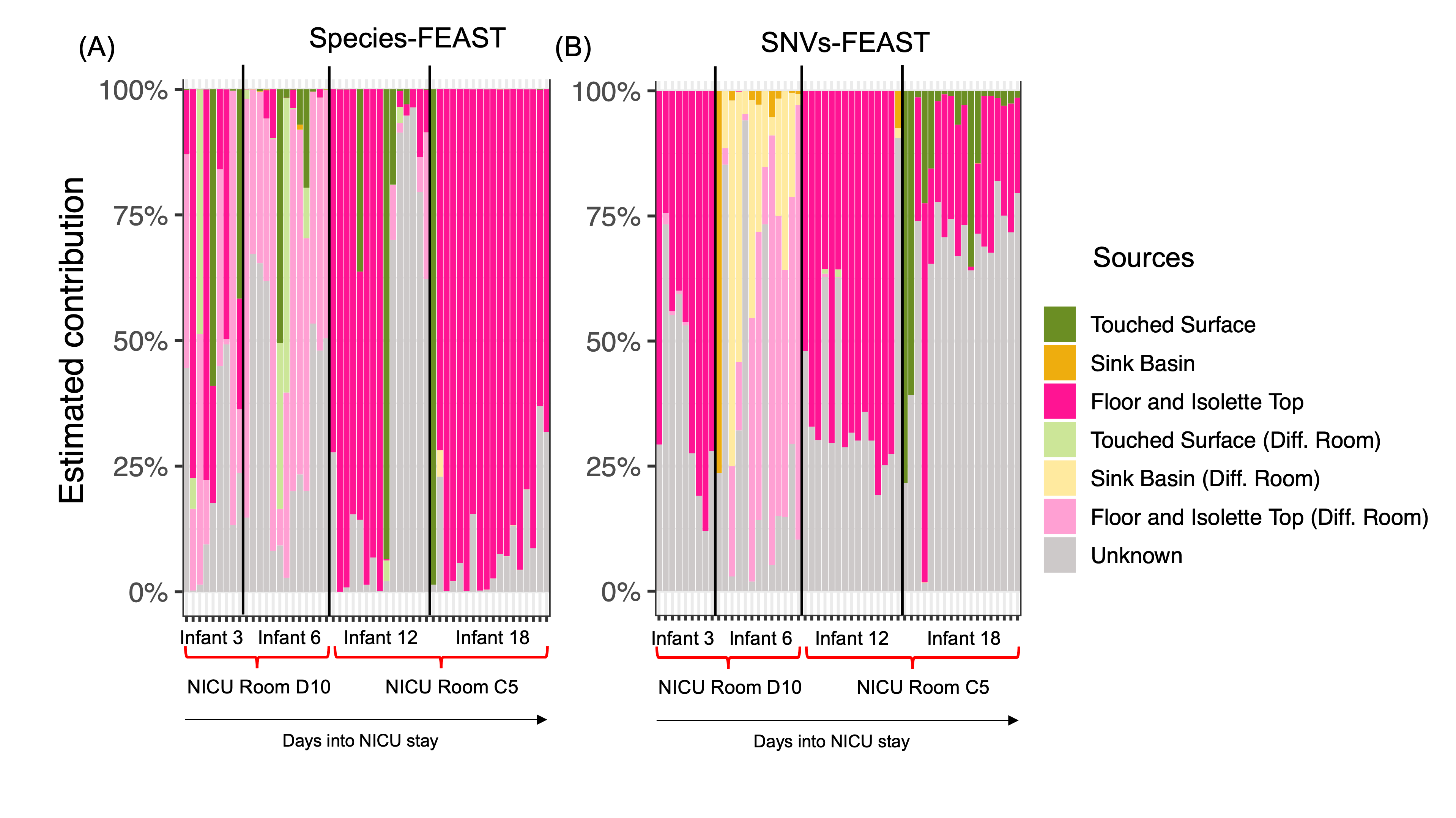

**Figure S6: Microbial source tracking with infants in the NICU and their built environment.** Contribution of samples from either the infant’s own NICU room or a different room from the study estimated using (A) species-FEAST and (B) SNV-FEAST. This is the same data that is plotted in **Figure 4A**, except all potential sources are stacked. This permits visualization of proportion unknown.

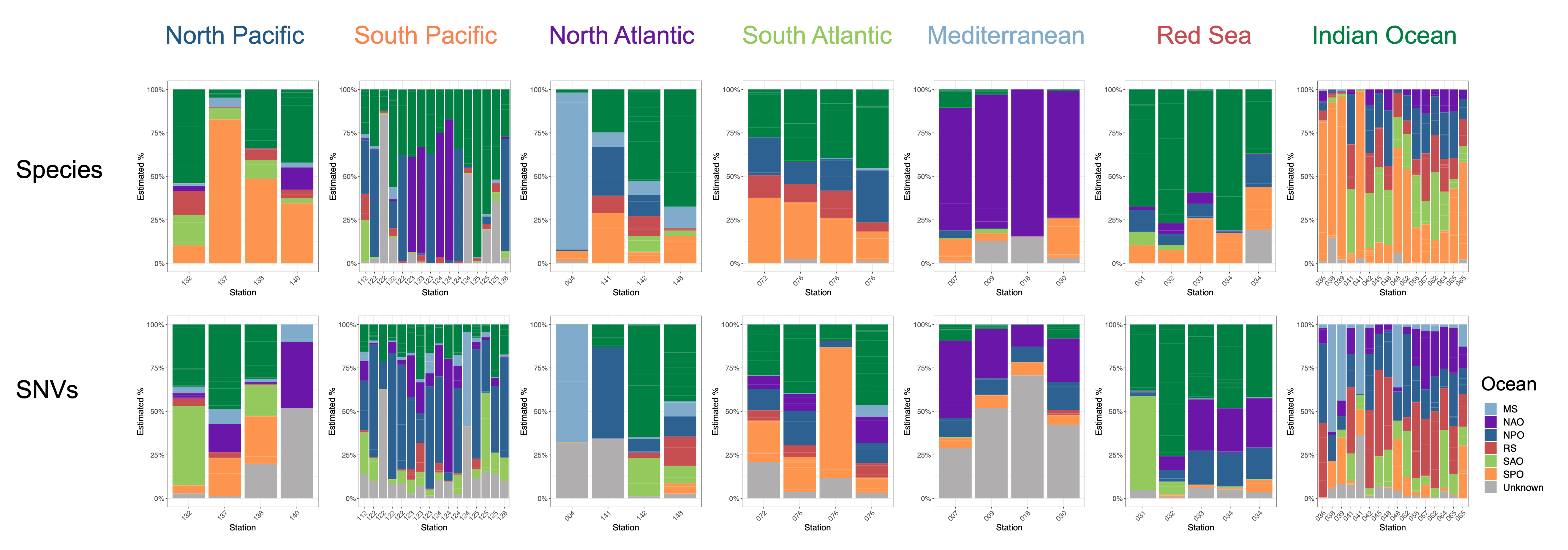

**Figure S7. Microbial source tracking in the Tara Oceans dataset with SNV and species-FEAST.** Source tracking estimates for the contribution of different oceans are depicted with vertical bars for the North Pacific (n=4), South Pacific (n=16), North Atlantic (n=4), South Atlantic (n=4), Mediterranean (n=4), Red Sea (n=5), and Indian Oceans (n=16). In each experiment, all stations around the world excluding those from the “sink” ocean are considered potential sources.

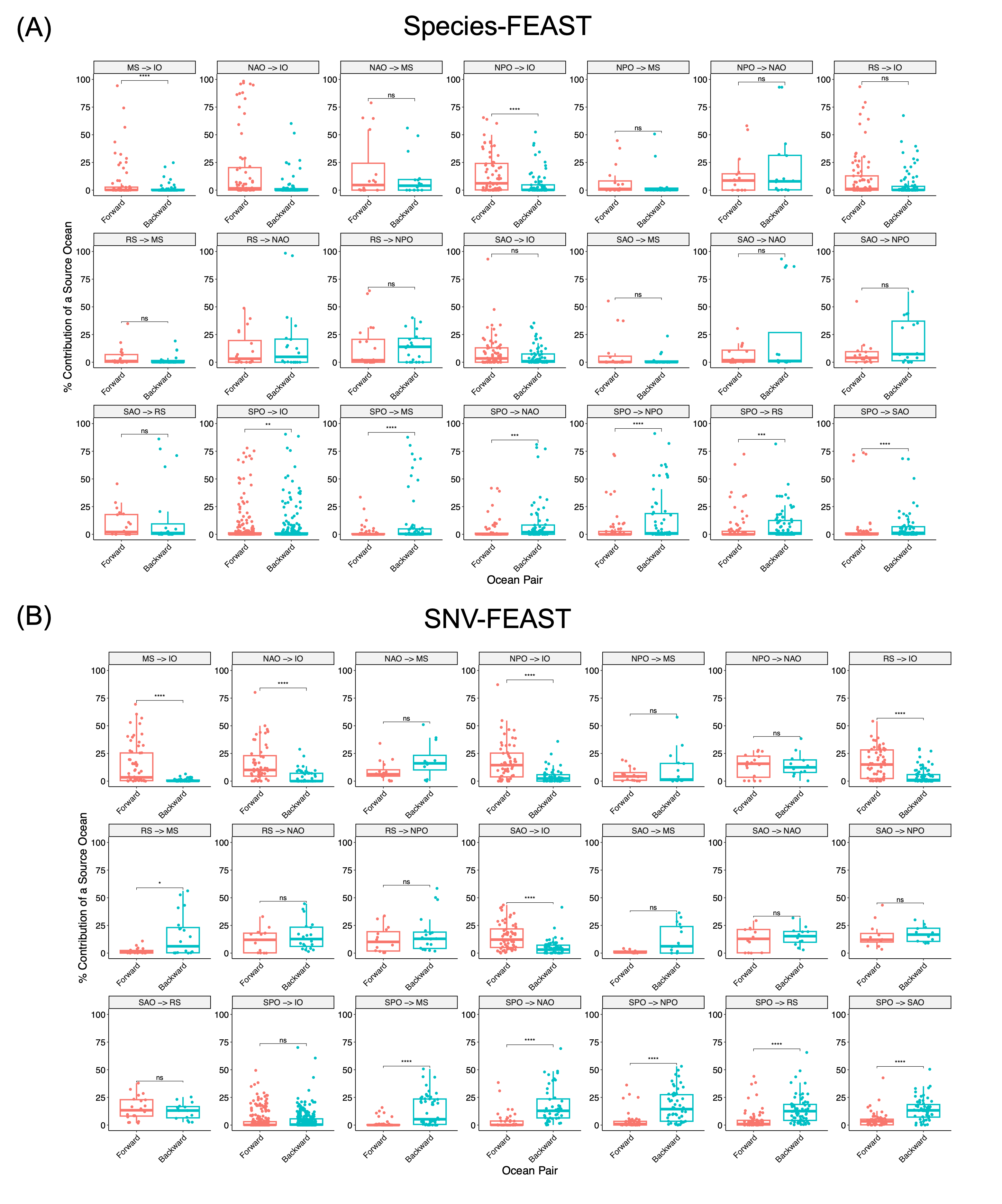

**Figure S8. Flipped source tracking for all ocean pairs** Shown are (A) species-FEAST and (B) SNV-FEAST estimates for contribution of one ocean to another. Each dot represents the contributions of each samples from the source ocean to the sink ocean of interest.
